# In search of nonlipogenic ABCA1 inducers (NLAI): precision coregulator TR-FRET identifies diverse signatures for LXR ligands

**DOI:** 10.1101/2025.09.12.675753

**Authors:** Megan S. Laham, Martha Ackerman-Berrier, Fahmida Alam, Sarah Turner, Ganga Reddy Velma, Christopher Penton, Soumya Reddy Musku, Manan Rana, Senthilkumar Thulasingam, Anandhan Annadurai, Maha Ibrahim Sulaiman, Nina Ma, Gregory R J Thatcher

## Abstract

APOE4, the major genetic risk factor for Alzheimer’s disease (AD), and ABCA1, required for lipidation of APOE are gene products of the liver X receptor (LXR) receptor. LXR agonists have been validated in animal models as therapeutics for AD, atherosclerosis, and many other diseases. Clinical progress has been thwarted by unwanted hepatic lipogenesis. Structurally diverse LXR ligands were profiled in coregulator TR-FRET (CRT) assays analyzing ligand-induced coactivator recruitment, coactivator selectivity, corepressor dissociation, and LXR isoform selectivity. A multiplex CRT assay was developed to measure synchronous ligand-induced displacement of corepressor by coactivator. Potency for coactivator recruitment to LXRβ correlated with induction of ABCA1 in human astrocytoma cells. Correlation with lipogenic activation of sterol response element (SRE) in hepatocarcinoma cells, was more complex. CRT response was diverse revealing ligands with theoretical full agonist, partial agonist, antagonist, and inverse agonist, and other signatures within the same chemical series, suggesting the scope for precision CRT to guide nonlipogenic LXR agonist design.

## Introduction

Definitive diagnosis of Alzheimer’s disease (AD) requires post-mortem hallmark neuropathology of amyloid-β (Aβ) plaques and neu-rofibrilliary tangles of intraneuronal tau protein, in combination with characteristic in-life cognitive decline. Alzheimer’s disease and related dementia (ADRD) defines a broader patient population, taking account of observations that: (i) hallmark pathology is not necessary and sufficient for progression to dementia; and (ii) at least half of AD patients manifest multiple forms of neuropathology at death. The original description, by Alois Alzheimer, of AD pathology in the brain of Auguste Deter, in addition to Aβ and tau, has been interpreted to include a third neuropathology: lipid granule accumulation in glial cells.^1^ Although this pathology, associated with lipid droplets, has been relatively forgotten, several modern theories of ADRD pathogenesis, such as the “small particle HDL (high density lipoprotein) hypothesis”^2^ propose a central role for cholesterol mobilization and lipoprotein transport. The 2020 Lancet Commission on Dementia stated that reducing the risk from diseases such as Type-2 diabetes (T2D) and obesity would have a significant impact on healthy aging with cognition intact.^3^ Subsequently, the 2024 Lancet Commission introduced LDL (low-density lipoprotein) cholesterol as the highest-contributing modifiable risk factor for dementia.^4-5^

Lipid droplets contain esterified cholesterol, phospholipids, triglycerides and other debris from phagocytosis of neurons and are found in astrocytes and microglia as a normal component of lipid homeostasis.^6^ The processing of dying cells, a primary task of microglia, relies upon cholesterol metabolism and transport, which when microglia are overwhelmed requires recruitment of astrocytes to assist in phagocytosis, leading to astrogliosis.^7-8^ Under physiological conditions, lipid droplets undergo lipophagy via mitochondrial cytochrome P450 (CYP)-mediated oxidation of cholesterol to 24-hydroxycholesterol (24HC) and 27-hydroxycholesterol (27HC), metabolites that cross the blood-brain barrier (BBB). Normal cholesterol metabolism and transport utilizes peripheral reverse cholesterol transport (RCT) for clearance to the liver, where cholesterol and metabolites are converted to bile acids. However, aberrant accumulation of lipid droplets in microglia and astrocytes leads to reduced lipophagy, accumulation of cholesterol and reduced production of 27HC. The consequence is elevated cholesterol levels in all cellular membranes, including the plasma membrane where lipid raft malfunction leads to disruption of synapses and multiple signaling pathways. Lipophagy and clearance of excess cholesterol via RCT requires efflux to lipoproteins, which is orchestrated by the nuclear hormone receptor (NR) liver-X-receptor (LXR) and its gene products, including *APOE* (apolipoprotein-E) and *ABCA1* (ATP-binding cassette-A1).^9^ *APOE4* is the major genetic risk factor for ADRD and is associated with aberrant lipid accumulation, notably in microglia.^10-11^

ABCA1 is expressed ubiquitously throughout the human body, with levels in the periphery highest in liver hepatocytes.^12^ The 2,261 amino acid human ABCA1 protein is an integral membrane protein consuming ATP to transport cholesterol, phospholipids, and other lipids to apolipoprotein carriers that bind to ABCA1.^13-15^ ABCA1 associates with cholesterol-rich membrane domains in the plasma membrane and intracellular organelle membranes, including endosomes and lysosomes.^16^ ABCA1 functions, in part, to protect cells from excessive and potentially cytotoxic accumulation of free cholesterol. ABCA1 transports lipids to specialized apolipoproteins, APOA1, APOA2, APOA4, APOC1-3, and APOE, to assemble HDL particles.^17^ ABCA1 is uniquely important in early loading of apolipoproteins in the minimally lipidated state,^18^ in contrast to related ABC family members (e.g. ABCG1/G4) and cholesterol transporters (e.g. SR-B1) that interact with apolipoproteins in their more lipidated states.^16, 19^ Owing to the direct influence of ABCA1 on cellular cholesterol homeostasis, increased ABCA1 activity has many indirect physiological functions. Taken together, the evidence supports ABCA1 as a promising target for therapeutics that might broadly attenuate pathophysiologic processes in the periphery and in the central nervous system (CNS). In the brain, ABCA1 is found in neurons, astrocytes, and microglia.^20^

ABCA1 associates with cholesterol-rich membrane domains, where it protects cells from excessive, and potentially cytotoxic, cholesterol and lipid accumulation.^13-16^ LDL particles contribute to vascular cholesterol deposition, inflammation, and atherogenesis to drive atherosclerosis.^21-23^ Conversely, HDL particles are anti-atherogenic. HDL facilitates removal of cholesterol from vascular walls and peripheral tissues to the liver for catabolism and excretion, via RCT. HDL, thereby prevents formation of cytotoxic oxidized lipid species and reduces inflammation and atherogenic lesion formation. ^24^ ABCA1 is essential for formation of nascent small HDL particles^14, 19, 25^, and with ABCG1, promotes formation of mature HDL particles.^26^

Reduced plasma cholesterol efflux capacity, which is directly tied to ABCA1 and ABCG1 expression, synergizes with other components of metabolic syndrome to promote atherogenesis.^27^ Oxidation of LDL and other pathological changes that occur during atherogenesis, may downregulate ABCA1 expression.^28-29^ Tangier disease, characterized by *ABCA1* loss-of-function mutations, leads to premature atherosclerosis;^30^ furthermore, *ABCA1* variants that increase AD risk have also been associated with T2D and cardiovascular disease (CVD) risk.^31-33^ Peripheral atheroprotection via ABCA1 induction promotes cerebrovascular health and preserves BBB integrity and function.^34-35^ Therefore, the beneficial role of ABCA1 in CNS disorders may derive from central actions and peripheral effects.^36-37^

*ABCA1* and other genes involved in cholesterol metabolism, mobilization, and signaling are gene products of LXR. Therapeutic approaches to enhance ABCA1 have been reviewed.^38-40^ LXR is the primary NR target for pharmacologic induction of ABCA1 expression. LXR agonists have been studied in many disease models, with the most intensive efforts targeting a therapeutic strategy to treat atherosclerosis and hypercholesterolemia.^41-45^

The two NR isoforms, LXRα (NR1H3) and LXRβ (NR1H2) mediate gene transcription via transcriptional complexes that bind to specific or shared NR response elements on DNA. LXRα is expressed in liver, small intestine, and adipose, and LXRβ is ubiquitously expressed. LXR forms heterodimers with retinoid X receptor (RXR). Transcriptional complexes are formed by recruitment of coregulator proteins that provide scaffolds for binding of epigenetic tools for DNA and histone writing, erasing, and reading. Since coregulators are shared between NRs, competition for coregulators is one of the many elements allowing crosstalk between different NRs. The canonical mechanism of ligand activation is the binding of an agonist to the NR dimer, in this case the LXR: RXR heterodimer, leading to coactivator recruitment to the transcriptional complex. In an alternate mechanism, transcriptional repression is maintained by corepressor binding to the unliganded heterodimer (**Fig. 1A**), with ligand binding leading to activation/derepression (**Fig. 1B**). The endogenous ligands for LXR are oxysterols (including 22(*R*)-hydroxycholesterol, 24HC, and 27HC) ^9, 46-47^ that activate LXR via canonical or alternate mechanisms.

**Fig. 1.**
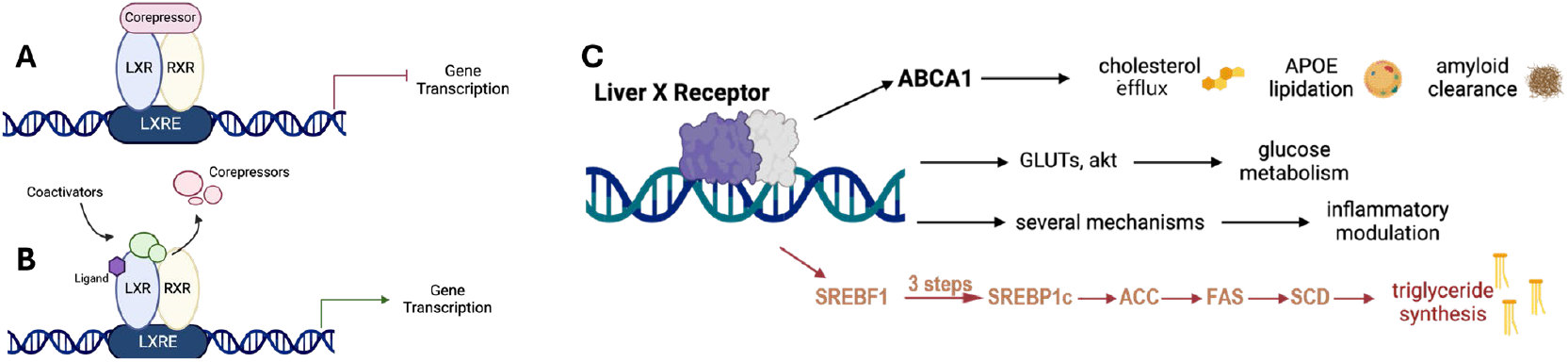
LXR activation mediates beneficial effects and unwanted lipogenesis. LXR transcriptional complexes are RXR heterodimers that can be activated by a canonical mechanism of coregulator recruitment or an alternative de-repression mechanism of (**A**) a constitutively repressed complex by (**B**) corepressor displacement. (**C**) ABCA1 gene transcription leads to cholesterol efflux and mobilization, APOE lipidation and potentially Aβ clearance, in addition to other mechanisms of potential benefit in ADRD; whereas, SREBF1 transcription initiates a lipogenic program in the liver leading to unwanted LDL formation and TG elevation.

LXR controls several transcriptional programs one of which, mediated by the master hepatic regulator sterol-response element binding protein 1c (SREBP1c) leads to unwanted lipogenesis, triglyceride (TG) synthesis, and elevated LDL (**Fig. 1C**). Adverse effects associated with elevated levels of TG, include steatohepatitis, and hepatomegaly.^7^ The challenge in developing LXR agonists as therapeutics for T2D, CVD, and ADRD has been to avoid the adverse activation of the lipogenic axis (**Fig. 1C**). The dominant approach has been to develop agonists selective for LXRβ over LXRα.^8^ The rationale for developing LXRβ-selective agonists is based on the high expression of LXRα in the liver; however, despite considerable efforts by pharmaceutical companies, selective LXRβ-selective agonists have yet to be successfully translated to the clinic.^48-49^

A recent paper from Astra Zeneca researchers (Belorusova et al. ^50^) specifically addressed the challenge in discovery of LXR ligands as nonlipogenic ABCA1 inducers to treat atherosclerosis. Mouse intestinal ABCA1 and plasma TG were used as binary ligand descriptors that were correlated with LXR readouts, most notably H/D-exchange mass spectrometry (HDX-MS) measurements that inferred ligand-induced conformational changes in the LXR ligand binding domain (LBD).^50^ Coregulator TR-FRET (CRT) measurements of coregulator binding to the LBD have been widely used in the NR field.^51^ A detailed and comprehensive CRT study of LXR ligands, both clinical and late preclinical, has not been reported.^52^ CRT data has been reported for individual ligands; for example, to claim that LXR agonist IMB-808 was nonlipogenic because of selective recruitment of specific coactivators. ^53-54^ Since repression of lipogenesis in the liver requires corepressor binding to LXRα, corepressors must be included in CRT studies.

We profiled diverse LXR ligands by CRT, including those studied in clinical trials: (i) to assess NR promiscuity by measuring coactivator recruitment to 9 steroid NRs; (ii) to measure LXRα and LXRβ selectivity by measuring SRC1 coactivator recruitment; (iii) to measure differential recruitment of other coactivators; (iv) to delineate the effect of putative antagonists, partial agonists, and inverse agonists; and (v) to define corepressor binding in the presence and absence of a coactivator using multiplexed precision CRT (pCRT).

CRT data was correlated with data from reporter assays in astrocytoma and hepatocarcinoma cells, measuring induction of ABCA1 and SREBP1c, respectively. The results from CRT analysis reveal a surprisingly diverse set of responses that would support diverse phenotypes. NRs are viewed as druggable targets given that the efficacy of 16% of approved drugs is mediated by NRs.^55^ The clinical success of selective estrogen receptor modulators (SERMs) is based upon tissue-selective transcriptional activity that is not predictable *a priori* but is thought to be governed by coregulator recruitment.^56^ More precise understanding of ligand-specific coactivator and corepressor binding will provide a predictive framework for nonlipogenic LXR modulators and reinvigorate the therapeutic targeting of NRs by small molecules.

## Results and Discussion

### Selection of benchmark LXR ligands

First reported in 2000^57^, T0901317 (T0) is commonly used as a benchmark or control LXR agonist. Cross-reactivity of LXR agonists with other NRs has been reported.^58^ T0 has been claimed to be an agonist for farnesoid X receptor (FXR) and pregnane X receptor (PXR) ^59-61^; although, in our CRT assays potent PXR agonism was observed without evidence for FXR binding. The cross-reactivity of T0 with PXR could confound its use as a control LXR agonist in cellular systems. However, T0 is unique in showing no selectivity between LXR isoforms, which makes it ideal for normalization of CRT data. T0 is the most widely studied LXR agonist, often as a control compound or LXR chemical probe, with dozens of papers reporting effects in preclinical animal models of disease, including AD.^62-65^ T0 is reliably lipogenic in mouse, hamster, and other models, causing elevated TGs and hepatic steatosis.

**Scheme 1.**
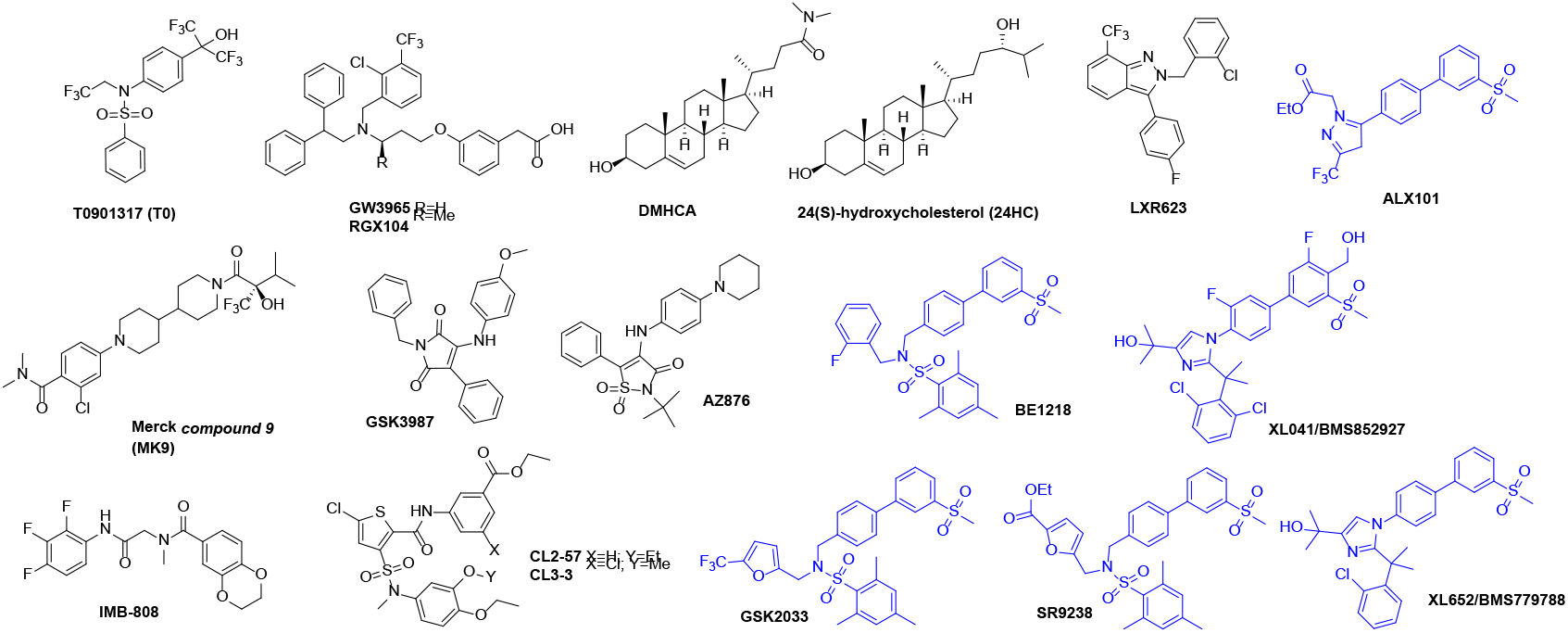
Chemical structures and compound codes of literature LXR ligands with biphenyl sulfone ligands shown in blue.

A different chemotype was identified using a cell-based ABCA1 reporter system, leading to GSK3987, an LXR agonist that in cell cultures showed concentration-dependent increases in ABCA1, SREBP1c, and cholesterol efflux and concentration-dependent decreases in IL-6.^66^ There is a lack of *in vivo* data, which may explain why this compound has not been used as a chemical probe.

GW3965, first reported in 2002, is widely used as an orally bioavailable selective LXR agonist in preclinical animal models. ^67-68^ Although the earliest report on GW3965 indicated PXR agonist activity, no binding to PXR is seen in CRT (*vide infra*). In C57Bl/6 mice, the effects of T0 and GW3965 were compared after 3 days treatment, measuring plasma lipids and gene induction in the liver and small intestine.^68^ Responses were highly dose-dependent; however, T0 caused increases in liver weight and TGs, whereas even at 100mg/kg/day, GW3965 had no effect on these liver markers. T0 induced SREBP1c and greatly induced fatty acid synthase (FAS) (38-fold) while GW3965 induced minimal increases in SREBP1c and FAS expression in the liver. GW3965 administration resulted in ABCA1 expression increased 8-fold in the small intestine and 7-fold in peripheral macrophages. Recent dramatic evidence of the benefits of ABCA1 induction by GW3965 was reported in a tauopathy mouse model relevant to AD.^69^ The beneficial effects of low dose GW3965 were replicated by genetic upregulation of *ABCA1*.

The LXR agonist, RGX104 (Abequolixron or SB742881), described as a first-in-class immunotherapy is being studied in cancer clinical trials. RGX104 is the methyl derivative of GW3965 and was compared to GW3965 in hamsters.^70^ Hamsters, unlike mice, express cholesteryl ester transfer protein (CETP) and are argued to have a more humanlike lipid metabolism. Both GW3965 and RGX104 caused a dose-dependent increase in plasma TG, LDL-C, and very-low-density-lipoprotein cholesterol (VLDL-C). Despite structural similarity, RGX104 significantly increased liver triglycerides but GW3965 did not. Similarly, hepatic gene transcription was weaker for GW3965, although comparable in intestines and peritoneal macrophages. In cynomolgus monkeys, GW3965 increased LDL and HDL at 7 days treatment only, with no effect on TG; whereas, RGX104 gave a robust time-dependent increase in TG and LDL. Without overinterpretation, this study hints at possible tissue-selective actions differentiating GW3965 from RGX104: tissue selectivity is well-known for modulators of other NRs.

Cholesterol mobilization was compared in HepG2 cells for RGX104, GW3965 and T0.^71^ In these cell cultures, all LXR agonists gave similar induction of ABCA1, ABCG1, FAS, and SREBP1c, which was argued to result in serum-dependent cholesterol efflux with induction of compensatory cholesterol synthesis at a level still leading to cellular cholesterol depletion. Interestingly, steroidal agonists, 22(R)-hydroxycholesterol, 24(S),25-epoxycholesterol, and DMHCA (N,N-dimethyl-3β-hydroxy-cholenamide) did not induce the same response.

The synthetic steroidal LXR agonist, DMHCA, was reported to increase LXRα, ABCA1 and SREBP1c genes in HepG2 and THP-1 cells to a lesser extent than T0 but to decrease SREBP1c in the J774 macrophage cell line and not significantly elevate FAS in THP-1 cells.^72^ In THP-1 cells, DMHCA attenuated SREBP1c gene transcription elicited by GW3965 co-treatment, implying partial agonist activity. C57BL/6 mice treated for 6 or 7 days with T0, GW3965, or DMHCA showed increased hepatic gene expression of ABCA1, ABCG1, SREBP1c, FAS, and acetyl-CoA carboxylase (ACC) for all agonists. In mice, T0 and GW3965 were reported to increase plasma TG and VLDL-C. DMHCA is metabolically labile with poor oral bioavailability^72^; nevertheless, low dose oral delivery has been studied in the ob/ob mouse in the context of diabetic retinopathy and bone marrow pathology.^73^

Like DMHCA, LXR-623 (WAY-252623) is referred to as an LXR partial agonist.^74^ Comparison of LXR-623 with GW3965 in LDLR^-/-^ mice confirmed the expected anti-atherosclerotic efficacy of both LXR agonists.^75^ In hamsters, LXR-623 did not cause a dose-dependent increase in plasma TG. In primates a significant increase in liver TG was observed but only at high dose (50 mg/kg/day p.o.) and returned to baseline at day 7 of treatment. Tissues from primates showed elevation of *ABCA1* and *ABCG1* in duodenum and suppression of hepatic genes including *SREBF1, FAS, CYP7A1*, and *LDLR*. The authors concluded that unfavorable lipogenesis was not a class effect of LXR agonists. In the only report on the LXR-623 clinical trial, no data on lipogenesis is reported. Detailed PK/PD correlations were made with ABCA1 transcripts in blood samples.^76^ No serious adverse events were reported; however, all 6 participants on the highest 300 mg dose reported neurologic or psychiatric adverse effects of unknown cause.

In contrast to LXR-623, the clinical trial of XL041 (BMS85297) reported on both activation of ABCA1 and lipogenic genes. As seen for LXR-623, treatment with XL041 increased ABCA1 expression in trial subjects.^48-49^ XL041 demonstrated good pharmacokinetics, safety, and lipid profiles in mice and cynomolgus monkeys ^49^. However, in clinical trials, after multi-day dosing, elevated plasma and liver lipids and neutropenia were observed in healthy and statin-treated subjects. Notably, co-administration of statins caused incomplete attenuation of lipid elevation. ^49, 77^ The peripheral side effects observed with XL041, included neutropenia and elevated TG and LDL levels, both of which were reversible. No serious adverse effects were reported.

Published data from Merck’s LXR program is limited. *Compound 9* (MK9) was studied in 12-month-old Tg2576 familial Alzheimer’s disease (FAD) mice dosed subcutaneously for 3 weeks at 50 mg/kg/day.^78^ Significant increases in ABCA1 and apoE protein levels in the brain were not accompanied by significant changes in plasma or liver TG, in contrast to mice treated with T0. In monkeys administered MK9 (20 mg/kg qd p.o.) for 2 weeks, in cerebrospinal fluid (CSF) a threefold increase in APOE and Aβ was observed: CSF Aβ is often associated with enhanced clearance from the brain. These changes were accompanied by no change in liver fat content relative to control, in contrast to T0.

AZ876, classified by Belorusova et al. as lipogenic in mice,^50^ showed dose-dependent elevation of plasma TG accompanied by a 91% reduction in atherosclerotic lesion area. ^79^ Rescue of cardiac hypertrophy in mice using low dose AZ876 was not accompanied by increases in plasma TG. ^80^

CL2-57 was the result of early optimization of a hit discovered from cell-based reporter screening for activation of ABCA1-luciferase in astrocytoma cells, counterscreening using a SREBP1c-luciferase reporter in HepG2 cells.^81^ In an obesogenic high-fat diet (HFD) mouse model, CL2-57 corrected T2D-associated readouts, attenuated diet-induced increases in plasma and hepatic TG levels and ameliorated other HFD-associated metabolomic changes.^82^ Further optimization guided by phenotypic screening resulted in CL3-3, a more potent LXR agonist and ABCA1 inducer. CL3-3 was tested in E3/4FAD mice that express one copy each of the ε-3 and ε-4 alleles of human APOE, mimicking the majority of AD patients. These mice also express five mutant amyloid precursor protein (APP) and PSEN1 human transgenes linked to FAD. Treatment with CL3-3 increased ABCA1 expression, enhanced APOE lipidation and reversed multiple AD phenotypes, without increasing TG. This study is the first in a human *APOE*-expressing model with hallmark amyloid-β pathology

ALX101 (Rovazolac), SR9238, GSK2033, XL652, and BE1218 are examples of the many reported LXR ligands in the biphenyl-3-methylsulfone class, which also includes XL041. Reported in a 2013 patent publication, ALX101 is listed as the subject of a 2018 clinical trial in moderate atopic dermatitis. XL652, an LXR agonist studied in the program that led to XL041, is described as an LXRβ-selective partial agonist.^83-84^ In mice XL652 administration caused increased ABCA1 and ABCG1 induction with elevated plasma triglycerides only at the highest dose (30 mg/kg qd), accompanied by induction of hepatic SREBP1c but not FAS.

The structurally related sulfones, BE1218, SR9238, SR9243, and GSK2033 have all been described as LXR inverse agonists. SR9238 was claimed amongst the first synthetic LXR inverse agonists to be discovered. SR9243 is systemically available; whereas, the metabolically labile SR9238, owing to the metabolism of the ester linker, is reported to show liver selectivity for suppression of lipogenic LXR gene products. ^85-87^ In obese mice on HFD, plasma TG was suppressed and in a mouse model of non-alcoholic steatohepatitis (NASH), SR9238 treatment significantly reduced hepatic inflammation and fibrosis. In cell-based models, the activity of GSK2033 was consistent with LXR inverse agonism; however, in a NASH mouse model, the expression of lipogenic genes, including FAS and SREBP1c, was *increased* rather than suppressed and no beneficial effects on hepatic steatosis were observed. Subsequently, GSK2033 was reported to activate PXR, FXR, and RXR, negating utility as an in vivo chemical probe.

IMB-808 is structurally distinct from most other LXR ligands. In cell-based assays, it was claimed to activate LXR with modest selectivity for the α-isoform and to be nonlipogenic in HepG2 cells, which is the rationale for inclusion of this compound in our study.^88^ The selection of other LXR ligands, introduced above, provides a diverse array of structures, selectivity and both lipogenic and nonlipogenic properties. Although most compounds discussed have demonstrated beneficial responses in preclinical models of human disease, the lack of a predictive algorithm for ligand design to avoid or circumvent unwanted lipogenesis has prevented progress in the clinic.

### Agonists, antagonists and NR coregulator TR-FRET (CRT)

The ligand binding pocket of NRs can be considered as both orthosteric and allosteric, because it binds endogenous agonists and induces a conformational change that induces recruitment of coregulator proteins to helix-12 (H12) of the NR and hence regulates the transcriptional function of the NR complex at its cognate DNA response element. The labels applied to individual NR ligands (full agonist, partial agonist, antagonist, inverse agonist) can be ambiguous, leading to misuse. Owing to the tissue and cell-specific output resulting from ligand binding, individual genes may be regulated in different directions.

In the case of estrogen receptor (ER) ligands, the term “pure antagonist” has been coined and is usually, but not universally, used to describe a ligand that competitively displaces estradiol and induces a conformational change incompatible with transcriptional activation.^89-90^ In the case of synthetic agonists (ligands that induce a response in the same direction as the endogenous agonist), the term “biased agonist” is useful for NR ligands and is related directly to tissue selectivity.^91^ Study of coregulator binding to the NR-LBD can potentially simplify the terminology used for ligand-dependent responses, which we will revisit below. However, first we introduce CRT to study cross-reactivity of LXR ligand binding to related NRs.

### CRT analysis of NR cross-reactivity

Class II NRs include: the NR1 subfamily, LXR, farnesoid X receptor (FXR), peroxisome proliferator activated receptor (PPAR), pregnane X receptor (PXR), retinoic acid receptor (RAR), constitutive androstane receptor (CAR); and the NR2 subfamily retinoid X receptor (RXR). Heterodimer formation of RXR with NR1 family members is a key step in formation of transcriptional complexes. Most LXR DNA binding sites are shared with RXR; however, only a fraction of RXR binding sites are shared with LXR and genome-wide binding profiles are quite different across cell and tissue types.^92^

Owing both to the permissive nature of the heterodimer complexes within the sterol NR family and to structural similarities between endogenous ligands and ligand binding pockets, LXR ligands may activate other NR complexes. Cross-reactivity of LXR agonists has been reported:^58^ T0 is claimed to be an FXR agonist;^59^ T0 and GW3965 are reported to be PXR agonists;^60-61^ and GSK2033 binds to FXR, PXR, CAR, RXR, and other NR LBDs in a cell-based assay.^93^ Therefore, all LXR ligands were examined in a Lanthascreen CRT assay measuring ligand-induced stabilization of the NR:coactivator complex and NR-specific ligand binding. All CRT screens of NR selectivity were performed with a terbium donor exciting a fluorescein-tagged coactivator as a measure of ligand binding and recruitment. Benchmarking used a known agonist for each NR.

NR-CRT assays were developed for RXRα, RXRβ, RARα, RARβ, FXR, PXR, and PPARδ, in addition to LXRα and LXRβ (**Table S1**). Assays for RXRα and RXRβ were optimized using the D22 coactivator peptide and the RXR ligands SR11237 and bexarotene as positive controls. Both benchmark RXR ligands were RXRα-selective full agonists with very similar potency and no response was observed for any of the tested LXR ligands (**Fig. S1A**).

FXR is critical for maintaining bile acid homeostasis that when disrupted leads to excessive accumulation of cholesterol and its actions are interleaved with those of LXR. FXR represses the expression of SREBP1c.^94^ LXR promotes fatty acid and triglyceride synthesis and storage, whereas FXR activation leads to decreased liver and plasma triglyceride levels.^95^ The CRT assay optimized for FXR used SRC2-2 as coactivator and obeticholic acid (OCA) as benchmark agonist, revealing no agonist activity by LXR ligands above baseline, despite description of T0 as a weak FXR agonist (**Fig. S1B**).^59-60^

PPAR isoforms have been reported to modulate ABCA1 expression levels. ^96-98^ PPARγ activity is closely associated with glucose metabolism, and PPARα and PPARδ activity is associated with fatty acid metabolism. PPARα and PPARγ agonists have been shown to activate ABCA1, improve TG levels, and have shown beneficial effects in FAD mouse models.^99^ PPARδ-selective agonists have also been reported to increase ABCA1,^100^ with examples in AD clinical trials.^101^ The CRT assay for PPARδ used a C33 peptide coactivator and GW501516 as positive control, with no activity observed for any LXR ligands tested (**Fig. S1C**).

Of the three isoforms, RARβ is the most widely expressed across-tissues: brain, intestine, liver, kidney, etc. ^102-104^ The RARα CRT was optimized with D22 coactivator peptide and RARβ with SRC2-2 coactivator peptide. TTNPB was used as the benchmark agonist for both RAR isoforms tested, possessing very similar and high potency for both isoforms. Maximal response was seen at 1 µM of TTNPB, a concentration at which no LXR ligand registered any response (**Fig. S1D**).

PXR is a xenobiotic nuclear receptor, responsible for upregulating metabolic enzymes for detoxification of exogenous and endogenous compounds and is highly expressed in the liver, intestine, and kidneys.^105^ The PXR LBD is larger than other sterol NRs and the hydrophobic ligand binding pocket reflects its function in accommodating a wide range of xenobiotics.^106^ Drugs that bind and activate PXR bear a risk of drug-drug interactions (DDIs).^107^ The bisphosphonate, SR12813, was used as a benchmark PXR agonist; however, T0 proved to be a full agonist and more potent (EC_50_ = 300 nM) than the benchmark (**Fig. S1E**).

### Activity and isoform selectivity of LXR ligands

The CRT selectivity assay incorporates GST-tagged LXR-LBD, anti-GST-terbium, biotin-SRC-1, and streptavidin-XL665. Notably, the 25 amino acid SRC-1 peptide is the only coactivator, amongst those tested for LXR binding, which has the fluorophore remote from the coactivator peptide: i.e., the only coactivator tested that uses a fluorophore labeled anti-tag antibody to bind the tagged coactivator rather than a fluorophore-labeled coactivator. Binding of an agonist to LXR induces recruitment of coactivator SRC-1 (steroid receptor coactivator 1; also known as nuclear receptor coactivator 1, NCOA1). When excited at 340 nm the terbium will emit at 620 nm exciting XL665 and emitting a signal reflecting coregulator binding to LXR. The assay measures the ligand-dependent EC_50_ for ligand-induced coactivator recruitment to LXR; the affinity of the ligand for the LXR:coregulator complex contributes to this potency. The strength of signal at saturating ligand concentration in CRT assays is usually interpreted as maximal efficacy. For sterol receptors, endogenous ligands, including 24HC, generally induce efficacy lower than synthetic ligands; thus, for LXR binding, T0 was used as a suitable control (**Fig. 2**).

**Fig. 2.**
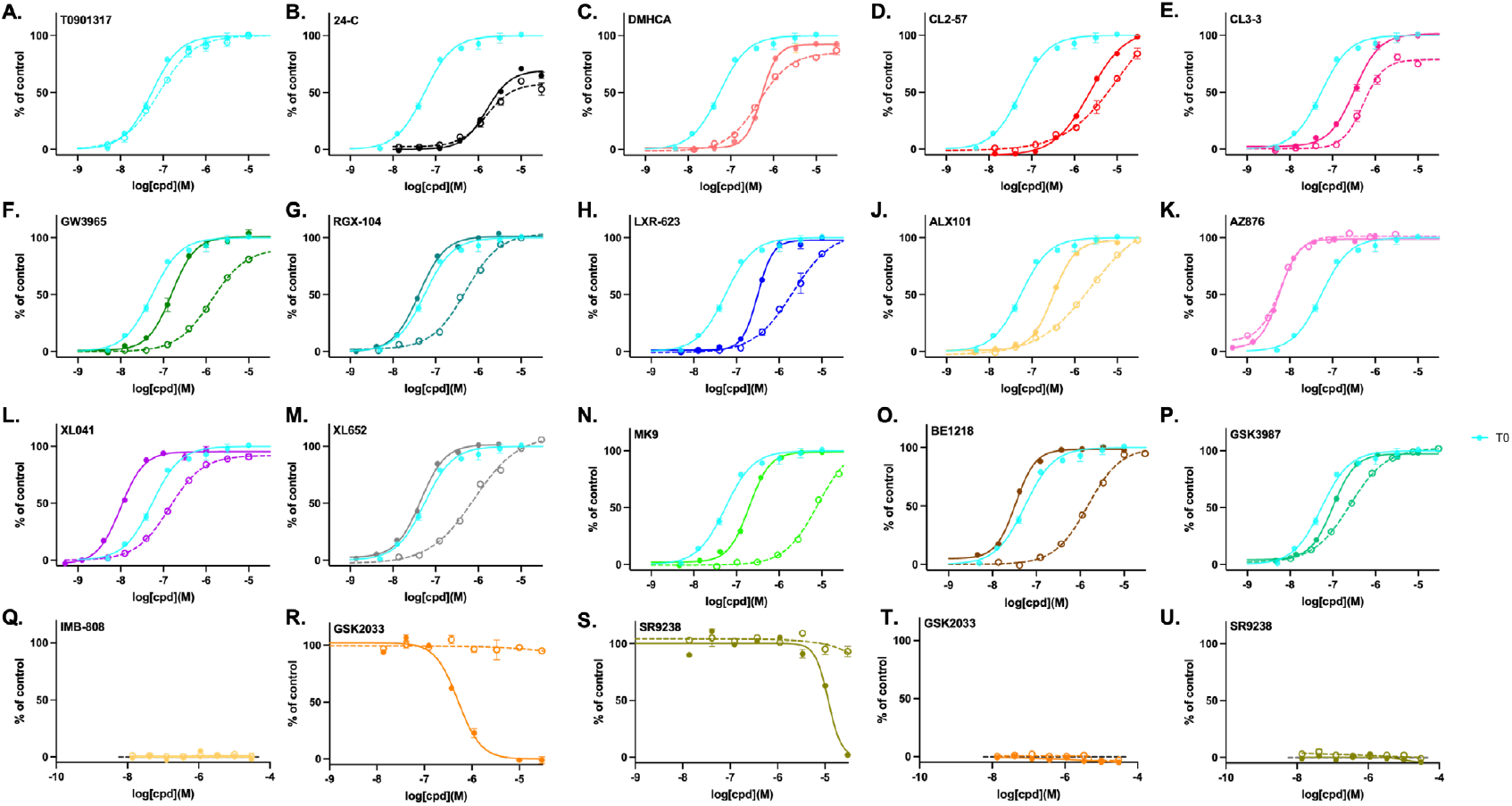
LXR isoform selectivity from CRT measurements. Concentration-response for recruitment of SRC-1 to LXRα (dotted lines) and LXRβ (solid line), determined by CRT assay, showing mean and SD from triplicate experiments, normalized to T0 (100%) as a full LXR agonist (shown in turquoise for reference). GSK2033 (**R**) and SR9238 (**S**) titrations were run in the presence of 24HC (3 µM) and neither ligand stabilized the LXR:SRC1 complex alone (**T**,**U**).

Although many LXR ligands have been reported to display partial agonist activity, most agonists studied in our biotin-SRC-1 CRT assay, gave maximal response at the same level as T0, behaving as full agonists. The clear exception is the endogenous partial agonist, 24HC; although, both the synthetic steroid DMHCA and CL3-3 showed partial agonist activity at LXRα. Both SR9238 and GSK2033 antagonized the agonism of 24HC at LXRβ without effect on LXRα.

In contrast to efficacy, both potency and selectivity were observed to vary widely. As expected, T0 was nonselective as were AZ876, DMHCA, and indeed 24HC. GSK3987 was weakly selective for LXRβ; whereas, most agonists displayed 10-100 fold selectivity for LXRβ over LXRα, with MK9 and BE1218 being the most selective LXRβ agonists. BE1218 was also one of the most potent LXRβ agonists; with the most potent being AZ876 (6 nM) and XL041 (10 nM). The most potent LXRα agonists were the pan-LXR agonists AZ876 (6 nM) and T0 (80 nM).

The ligand binding pockets of the two LXR isoforms differ by only one amino acid located in helix-3 (H3: LXRα-Val263 and LXRβ-Ile277). Interestingly, correction of this difference by mutation of these residues to alanine (V263A and I277A) was observed to reduce, but not to ablate isoform selectivity in reporter assays.^108^ Supported by modeling studies, this observation by Ding et al. led to the suggestion that conformational cooperativity of LBD residues beyond these two amino acids generally favors ligand binding to the LXRβ isoform. Therefore, most reported ligands, including those examined in the current work, are LXRβ-selective or non-selective.

### ABCA1 induction correlated with LXRβ not LXRα CRT potency

Astrocytoma CCF-STTG1 cells, stably transfected with an ABCA1-promoter linked to a luciferase response element, provide a model for ABCA1 induction in astrocytes.^81^ In contrast to the SRC1 CRT binding data, showing little evidence for partial agonist activity, concentration-response curves in cell-based reporter assays showed apparent partial agonist and antagonist activity (**Fig. 3**). The biphenyl sulfones, BE1218 and XL041, showed partial agonist response. The antagonist activity of GSK2033 and SR9238 in LXRβ:SRC1 CRT (**Fig. 1**) was reflected by inverse agonist activity in cells. Apparent inverse agonist activity in cell-based assays can result from “true” inverse agonist recruitment of corepressors or from antagonist activity towards endogenous sterol LXR agonists. Although we use serum-stripped media, antagonism of endogenous LXR agonists cannot be ruled out. As introduced above, BE1218, SR9238, SR9243, and GSK2033 have been described as LXR inverse agonists in the literature; however, in our hands, BE1218 acted as a full agonist in recruitment of SRC1 and a partial agonist of cellular ABCA1 induction.

**Fig. 3.**
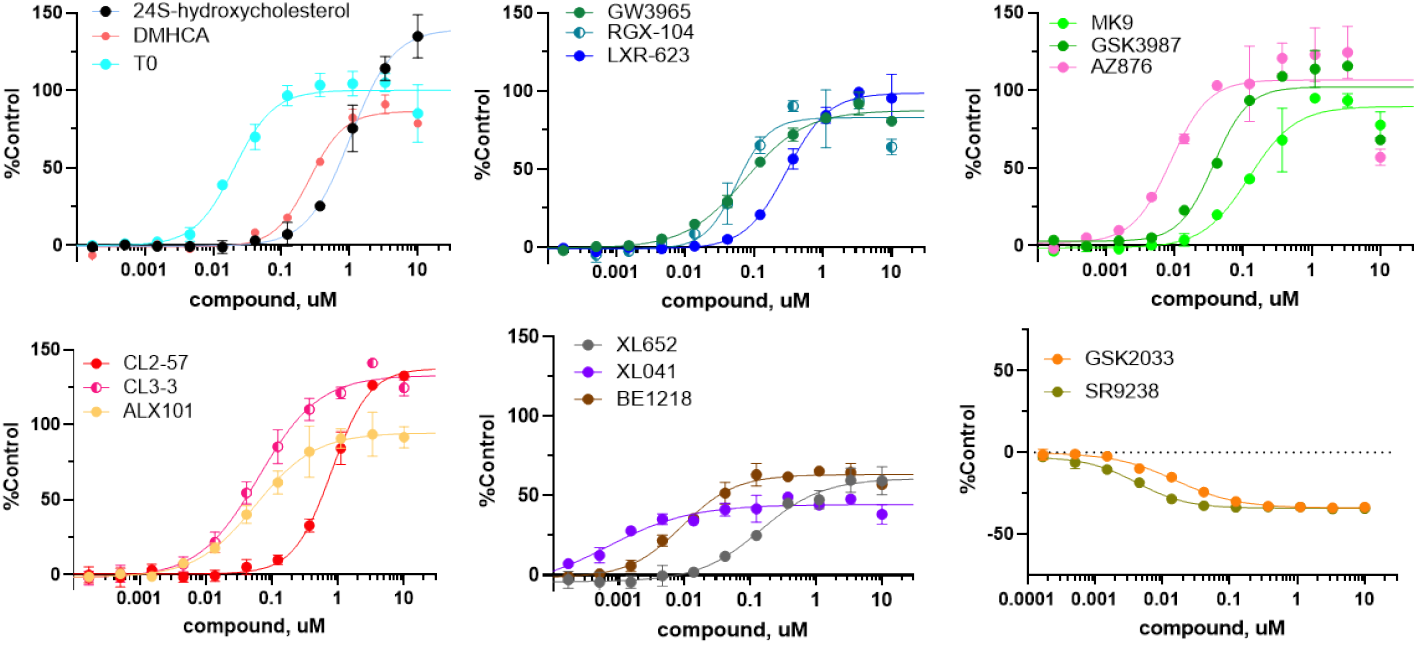
ABCA1-luc reporter in CCF cells. Concentration-response for LXR ligands in astrocytoma cells measured after 24 hr incubation, normalized to T0 response (100%) showing mean and SD from triplicate experiments (see **Fig. S2** for biological replicates).

The correlation of potency for cellular ABCA1 induction versus stabilization of the LXRβ:SRC1 coactivator complex was strong (Pearson r = 0.83 p = 0.0002; **Fig. 4A**); whereas, the correlation of ABCA1 induction with SRC1 recruitment to LXRα showed weak coupling (Pearson r = 0.6 p = 0.02; **Fig. 4B**). To the best of our knowledge, this is the first such quantitative ABCA1/LXR correlation across a varied structural array of LXR agonists and supports the targeting of LXRβ as a mechanism for induction of ABCA1 and related genes in cholesterol mobilization. The correlation between the CRT assays measuring potency for ligand-induced SRC1 binding to LXRα versus LXRβ was moderate (Pearson r = 0.7 p = 0.002; **Fig. 4C**), reflecting the presence of nonselective and LXRβ-selective ligands in the compound set.

**Fig. 4.**
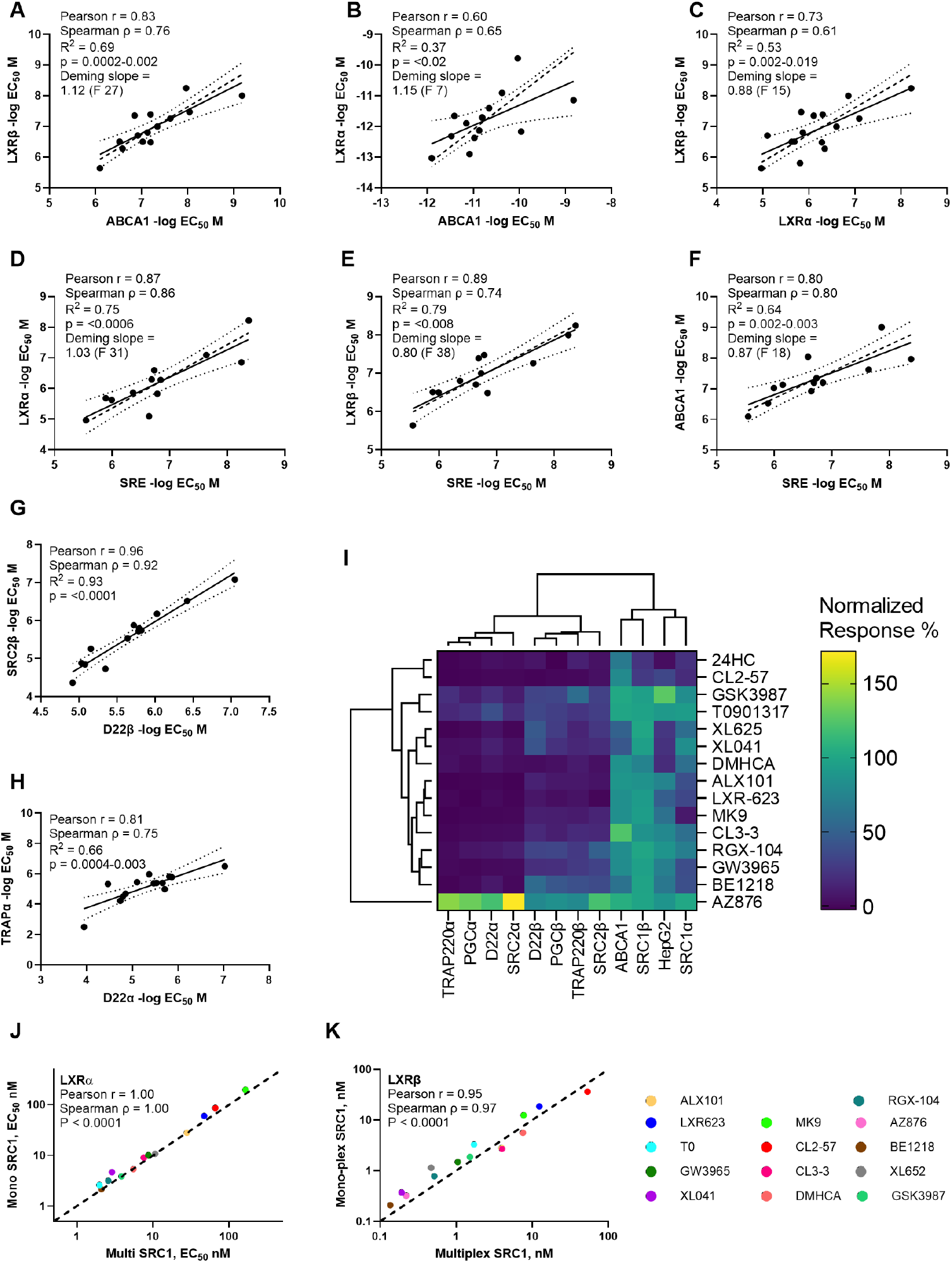
Correlating potency in cell-free and cell-based assays. **A-F)** Correlation of potency for SRC1 coactivator stabilization by LXR ligands from CRT data with potency for ABCA1 and SRE reporter assays in CCF and HepG2 cells, respectively. Pearson and Spearman correlation data are shown with significance and Deming slope with F-test, showing poor correlation of LXRα CRT data with ABCA1 activation. Solid line and 95% confidence limits are from simple linear fit with dashed line showing Deming fit. **G-H)** Correlation of potency for ligand-dependent recruitment of SRC2-2, D22, and TRAP220 to LXR α and β showing correlation statistics, best-fit line and 95% confidence intervals. **I)** Hierarchical clustering analysis of cell-free and cell-based relative response at 1 µM ligand. **J-K)** Correlation of potency for ligand-dependent recruitment of SRC1 to LXR α and β isoforms showing Pearson and Spearman correlations and best-fit line and 95% confidence intervals from simple linear regression.

### Lipogenic SRE activation and correlations with CRT data

Our phenotypic high throughput screen (HTS) approach to identify nonlipogenic ABCA1 Inducers (NLAI) originally used counterscreening with a SREBP1c-luciferase reporter (linked to the *SREBF1* gene promoter) in human hepatocarcinoma HepG2 cells.^81^ Unwanted lipogenesis is mediated by LXR agonists via direct transcriptional activation and indirectly by the effects of SREBP1c.^57 109^ SREBP1c enhances transcription of genes required for synthesis of fatty acids and TG and as the dominant isoform in liver is associated most closely with lipogenesis and TG elevation.^109^ SREBP1c is inactive and ER-resident, requiring transport by SREBP cleavage-activating protein (SCAP) to the Golgi membrane and proteolysis by S1P and S2P to release the mature SREBP1c transcription factor (**Fig. 1C**). Binding of SREBP1c to the sterol response element (SRE) induces transcription of lipogenic genes associated with elevated TG. To better model the lipogenic axis in hepatocytes, we use HepG2 cells stably transfected with a nanoluciferase reporter downstream of a SRE promoter (SRE-luc), activation of which requires SREBP1c maturation.

As observed with induction of ABCA1, concentration-response curves for SRE activation by LXR ligands showed apparent partial and full agonist responses (**Fig. 5**). At concentrations ≤ 1 µM, XL652 and XL041 activated SRE with low efficacy (20-25% of T0) and SR9238 lowered SRE activation below baseline. Three of the biphenyl sulfones, XL041, XL652, and SR9238, gave anomalous activation data at concentrations above 1 µM, associated with interference with the nanoluciferase reporter (**Fig. S3**). Owing to the anomalous response in the reporter assay, we measured lipogenic genes, *SREBF1, FAS*, and *SCD1* by RT-PCR in HepG2 cells in response to treatment with the biphenyl sulfones (**Figs 5G-I**). The PCR data was internally consistent, with GSK2033 and SR9238 reducing lipogenic gene transcription below baseline. Under the same conditions, GSK2033 and SR9238 showed no loss of cell viability relative to vehicle control (**Fig. S3**).

**Fig. 5.**
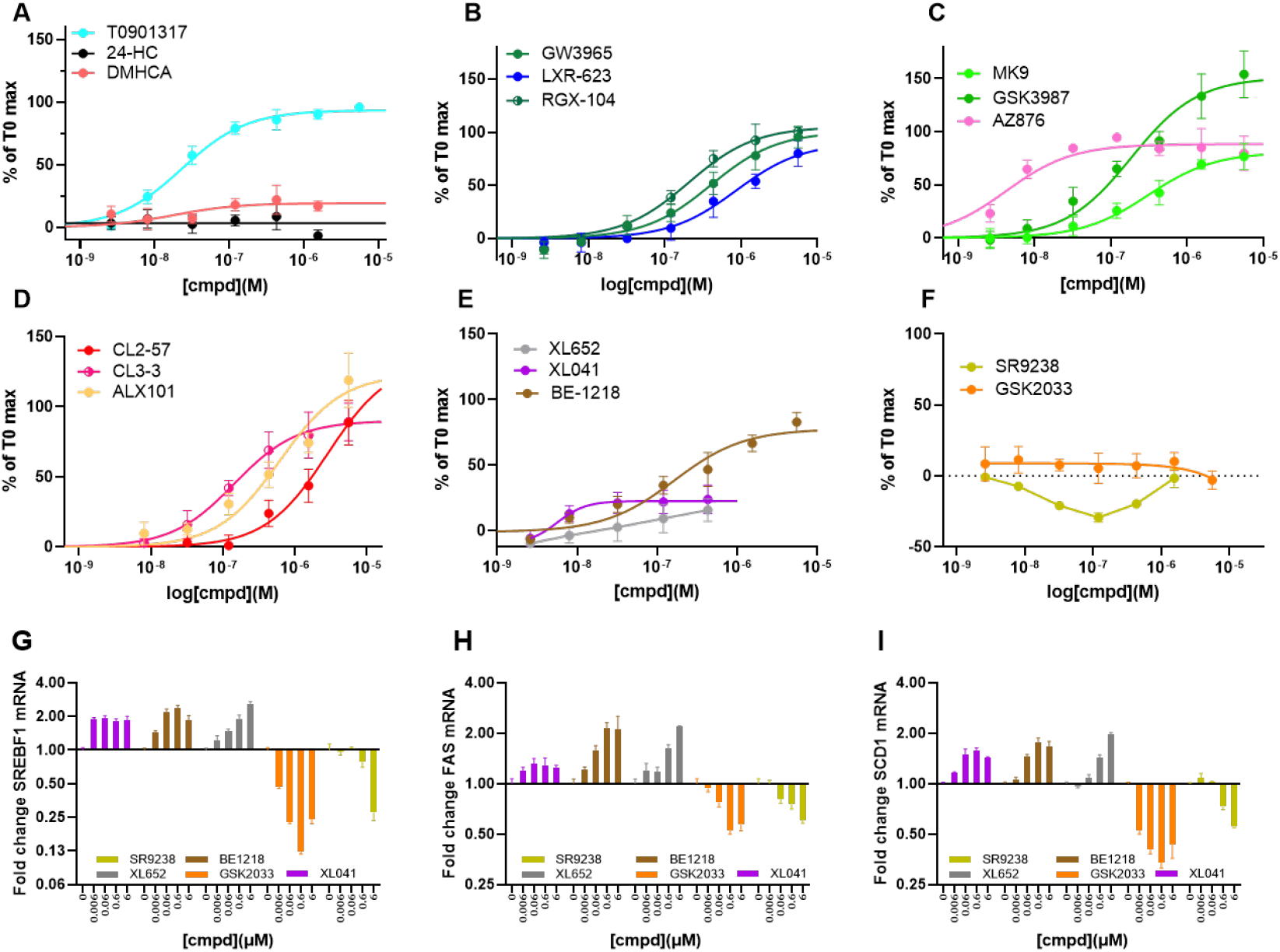
Lipogenic response of HepG2 cells to treatment with LXR ligands. (**A-D)** Concentration-response curves for LXR ligands normalized to maximal response for T0 (EC_50_ = 15 nM) showing a range of potency (AZ876 EC_50_ = 4.2 nM to CL2-57 2.8 µM) and maximal efficacy (BE-1218 80% to GSK3987 150%). See **Fig. S2** for biological replicates. **(E-F)** Three biphenyl sulfones gave anomalous activation of SRE at higher concentrations (**Fig. S3**): XL041 gave a partial agonist response at submicromolar concentrations (EC_50_ = 5.7 nM E_max_ = 22%), while SR9238 and GSK2033 had a neutral or antagonist response. (**G-I**) The transcriptional response to the five biphenyl sulfones was studied by RT-PCR under the same conditions as for SRE-luc measurements replicating the lipogenic induction observed in the reporter assays; both GSK2033 and SR9238 reduced response below baseline. Both DMHCA and 24HC were cytotoxic at higher concentrations. Data show mean and standard deviation for triplicate measurements. See **Fig. S3** for full concentration-response curves.

The corollary to targeting LXRβ as a mechanism for induction of ABCA1 is that LXRα activation causes lipogenesis and must be avoided. Simplistically, activation of the lipogenic axis (SRE activation) might be expected to correlate with SRC1 recruitment to LXRα; however, CRT correlations are compatible with a role for both LXR isoforms in lipogenesis. Correlations of SRE activation with SRC1 recruitment to LXRα (Pearson r = 0.87 p< 0.0006; **Fig. 4D**) and LXRβ (Pearson r = 0.89 p< 0.008; **Fig. 4E**) both gave strong and similar coupling. These correlations are consistent with the failure of LXRβ-selective ligands to circumvent lipogenesis in the liver.

A moderate correlation was observed between SRE activation and ABCA1 induction (Pearson r = 0.80 p = 0.002; **Fig. 4F**), with BE-1218 and XL041 outliers from the correlation with selectivity for ABCA1 induction. The correlation analyses used potency measurements both for SRC1 recruitment and for activation of luciferase reporters, which does not incorporate a contribution from the lower efficacy seen for partial agonists in cell-based assays. On the basis of potency alone: (i) cellular ABCA1 induction can be dissociated from induction of lipogenic genes for some ligands; and (ii) ABCA1 induction can be predicted by LXRβ recruitment of SRC1 measured by CRT. Data analysis to this point does not provide a framework for prediction of lipogenic potency or efficacy. Further, the lipogenic response in cell cultures is more complex and varied than ABCA1 induction.

### Coactivator selectivity

The varied physiological functions of a particular NR are controlled by recruitment of specific coregulators.^110-111^ Coactivator families include the p160 family (SRC-1 (NCoA-1), SRC-2 (GRIP1, TIF2, or NCoA-2), SRC-3 (p/CIP, RAC3, ACTR, AIB1, or TRAM-1)), PPAR*γ* coactivator 1 (PGC-1*α*, PGC-1*β*), and TRAP/DRIP complex. Thyroid hormone receptor-associated proteins (TRAPs) were first studied in the context of the thyroid hormone receptor.^112^ A highly conserved LXXLL motif in the p160 family of coactivators is necessary for association of coactivators to the ligand-bound receptor. ^113^ Two subunits of the TRAP complex, TRAP220 and TRAP100, contain LXXLL domains.

Selective recruitment of coactivators to LXR may provide a rationale for putative ligand selectivity for nonlipogenic gene induction. The lack of recruitment to LXRβ of D22 and TRAP220 was proposed to underlie nonlipogenic actions of IMB-808. Kim et al. explored the mechanism of LXR-mediated transcription of metabolic genes with a focus on coactivator complexes that lead to the modulation of specific target gene expression.^54^ Two coactivators were identified as proteins that interact with LXRα in a direct and ligand-dependent manner: TRAP220 and TRAP80. Examination of activation by T0 and GW3965 indicated that TRAP220 stimulates promoters for SREBP1c and ABCA1, whereas TRAP80 was selectively lipogenic. Silencing TRAP80 in mice followed by treatment with GW3965 or T0 was reported to prevent hepatic steatosis and reduce hepatic TG levels; whereas, RCT-related genes and the anti-inflammatory effects of the LXR ligands were unaffected. This observation provides a potential mechanism for selective repression of SREBP1c-mediated lipogenesis caused by LXR*α*.

CRT analysis of coactivator recruitment to LXR of PGC1α, SRC2-2, D22, and TRAP220 demonstrated that potency for LXR agonists was strongly correlated for LXRβ for all coactivator pairs (**Fig. 4G**; **Fig. S4A-D**) and for LXRα was strongly correlated between SRC2-2/D22 and PGC1α/TRAP220 pairs but more weakly between D22 and TRAP220 (**Fig. 4H**). Concentration-response curves, normalized to maximal response for T0, are shown for D22 and TRAP220 recruitment (**Fig. 6**) and for all coactivators (**Fig. S5**). Across the four coactivators studied, T0 was nonselective both for LXR isoform and for coactivator identity; whereas, most other agonists displayed both full and partial agonist profiles in contrast to our observations on SRC1 recruitment. Thus, across the four coactivators studied, most ligands showed LXRβ selectivity based on maximal efficacy, notably MK9, CL3-3, and LXR-623. Regarding LXR isoform selectivity, AZ876, does not show isoform selectivity for SRC1 recruitment but selectively recruits SRC2-2, D22, PGC1α and TRAP220 to LXRα. GSK2033 and SR9238 are unable to induce an LXR conformation compatible with recruitment of any coactivator studied. Several ligands induced very minimal recruitment of coactivator TRAP220 to LXRα, including XL041, XL652, BE1218, CL2-57, and CL3-3. Most ligands were selective for D22 over TRAP220 recruitment, notably XL041 and XL652.

**Fig. 6.**
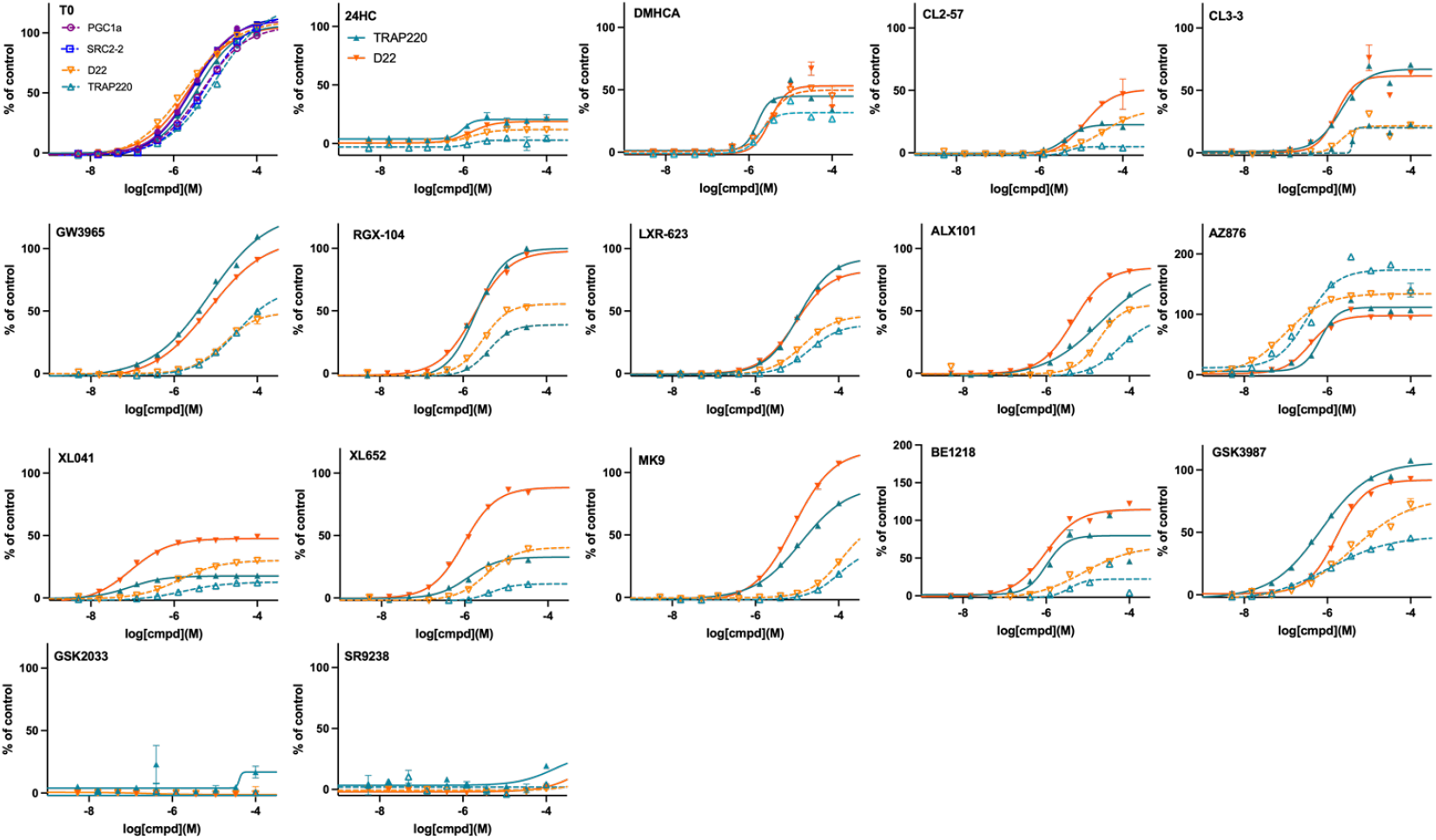
Coactivator recruitment to LXR induced by LXR ligands. CRT data was normalized to T0 maximal response (100%) for all four CoA studied: LXRα (dashed lines) LXRβ (solid lines). For clarity, only TRAP220 and D22 data are shown. As noted in the text, most ligands were β-selective or nonselective, with the exception of AZ876 that selectively stabilizes CoA binding to LXRα. GSK2033 and SR9238 were studied in the absence of agonist. Data shown mean and SD from at least triplicate measurements. (See **Fig. S5** all four CoA studied).

To reflect the contribution of potency and efficacy, relative response at 1 µM ligand was assessed by hierarchical clustering analysis for all ligands with agonist activity in CRT and reporter assays (**Fig. 4I**). Across ligands, AZ876 was differentiated as its own cluster owing to LXRα-selective activity in CRT assays; although, removing AZ876 from the analysis expanded clustering to 7 ligand clusters (**Fig. S4 G**,**L**,**M**). The dendrogram for assay response identified 4 clusters: 1) CoA/LXRα recruitment; 2) CoA/LXRβ recruitment; 3) ABCA1 activation with SRC1/LXRβ recruitment; and 4) SRE activation with SRC1/LXRα recruitment (**Figs 4I, S4 E**,**F**,**I**,**K**). These clusters fit with expectations based on linear correlations. A larger ligand set will enhance confidence in such clustering analysis.

### Corepressor stabilization and displacement

Support for the de-repression and canonical mechanisms of LXR activation (**Fig. 1**) comes from studies on selective knockout or dual knockout of LXR isoforms in mice and cells. As argued below, stabilization of corepressor binding to LXRα may bias towards nonlipogenic LXR agonist activity.

Liver-specific knockout of LXRα was reported to attenuate the effect of T0 treatment on elevation of lipogenic genes and plasma TG.^114^ Although T0-induced elevation of liver ABCA1 was also attenuated by LXRα^-/-^, these observations support the pursuit of LXRβ-selective agonists to minimize adverse effects in the liver. In LXRα^-/-^ mouse livers, the basal expression of some genes (*ABCA1* and *SREBF1*) was unchanged; whereas, others (e.g. *FAS*) were significantly decreased and *LXRβ* expression was increased.^13^ The elevation of *ABCA1, SREBF1*, and *FAS* induced by LXR agonists was attenuated in LXRα^-/-^ relative to wild-type (WT). In the livers of dual LXRα^-/-^/LXRβ^-/-^ mice, liver lipogenesis and TG levels were reported as lower with normal or high-cholesterol diets.^115^ In another study, increases in hepatic LXR target genes induced by T0 and GW3965 were attenuated in LXRα^-/-^ versus WT mice; however, higher doses of LXR agonists significantly elevated *ABCA1* and *SREBF1* in both mice.^116^ The observation that agonist-induced *ABCA1* expression was amplified in the intestines was interpreted to reflect the tissue selectivity of LXR agonists and desirability of selecting against LXRα agonism in the liver.

Transcriptomic analysis of LXR isoform knockouts adds complexity. In contrast to ubiquitous LXRβ expression, LXRα is localized in specific cells and tissues: e.g. liver and macrophages. Immortalized macrophages, LXR-null, or expressing either LXRβ or LXRα, were studied using GW3965 and/or GSK2033 as pharmacologic probes.^117^ Three distinct transcriptional mechanisms were proposed, one of which being pharmacologically nonresponsive and repressed in LXR-null macrophages can be assumed less relevant to LXR ligand design. In the classical or canonical mechanism (Mode II), transcription is weakly active or inactive in LXR-null and is active in the presence of an LXR agonist that induces a conformational change leading to coactivator binding. In the de-repression mechanism (Mode I), LXR-null is active, basal activity is repressed and the LXR agonist induces a conformational change that displaces the corepressor in favor of coactivator binding. *ABCA1* and *SREBF1* are used as examples of genes controlled by Mode I and Mode II mechanisms, respectively.

A similar transcriptomic approach to study of LXR-null mouse livers emphasized that knockout of LXR has significant effects on other nuclear receptors and transcription factors, notably PPARα.^118^ Perhaps surprisingly, the majority of genes differentially expressed in WT versus LXR-null liver were insensitive to treatment with T0 and GW3965. Genes downregulated in LXR-null and induced by agonists in WT livers included lipogenic genes *SREBF1, SCD1, FAS*, and *LPCAT3*. Although genes mediating cholesterol metabolism and efflux (e.g. *ABCA1, APOA1*) are induced by LXR agonists, these were generally unchanged in WT and LXR-null livers. Thus, in the liver, induction of *ABCA1* and *SREBF1* may be dominated by canonical and de-repression mechanisms, respectively.

Arguing that stabilization of corepressor binding to LXRα biases towards nonlipogenic activity, we used CRT to measure ligand-dependent, LXR:corepressor interactions. Nuclear receptor corepressor (NCoR) and silencing mediator of retinoic acid and thyroid hormone receptor (SMRT) are large proteins (ca. 270 kDa) that share about 40% sequence identity. SMRT and NCoR are largely disordered platform proteins that act as a scaffold upon which the enzymatic machinery of the repression complex is built.^119^ In a transcriptionally repressed complex, a corepressor typically binds to the LXR LBD.

The interaction motifs of coactivator and corepressor proteins compete for LBD binding. Binding of the coactivator interaction motif LXXLL is dependent on hydrophobic amino-acids in helix 12.^120^ The corepressor interaction motif (φxxφφ where φ is a hydrophobic amino-acid and x is any amino acid) uses additional flanking sequences, but contains a similar amphipathic core to the coactivator motif.^121-123^

NCOR2 and SMRT2 were observed to bind to LXR isoforms in the absence of ligand, with CRT analysis showing the expected displacement of corepressor (loss of signal) by LXR agonists (**Fig. 7**). For some ligands, stabilization of the corepressor-bound complex (gain of signal) was observed (**Figs 7, S6**). Titration with LXR agonists, GW3965, its methyl-derivative (RGX-104), and T0 caused displacement of NCOR2 from both LXR isoforms; whereas LXR-623 and DMHCA had little or no effect (**Fig. 7A,B**). A similar pattern of ligand response was seen for binding of SMRT2 to LXR (**Fig. 7C,D**). Across isoforms and corepressors, several ligands showed little or no ability to displace NCOR2 and SMRT2 corepressors from LXR: CL2-57, CL3-3, LXR-623, and the antagonist GSK2033 (**Figs 7, S6**).

**Fig. 7.**
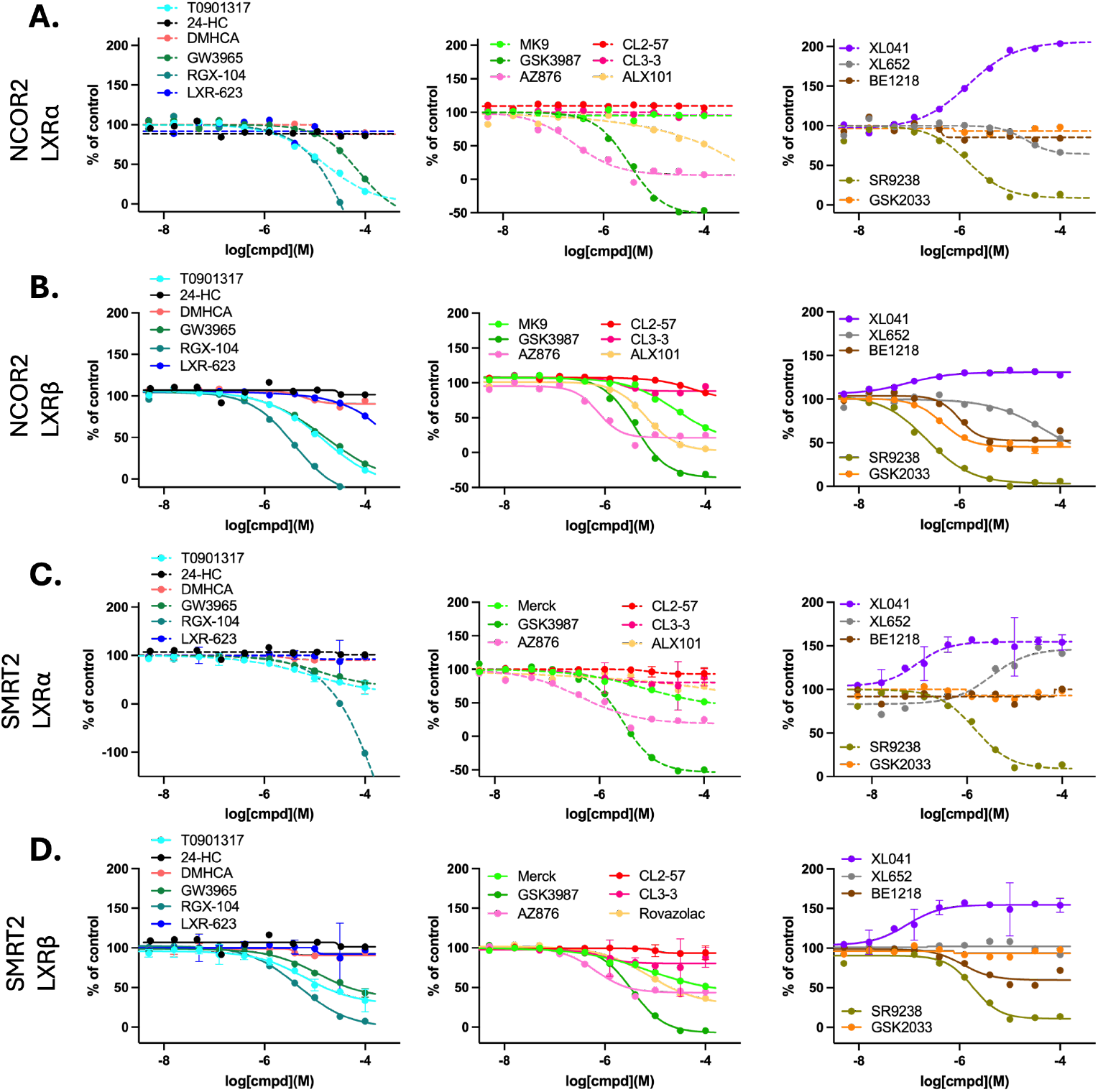
Corepressor binding to LXR isoforms: ligand dependence. Concentration-response from CRT measurements of: NCOR2 binding to LXRα (dashed lines) (**A**); NCOR2 binding to LXRβ (solid lines) (**B**); SMRT2 binding to LXRα (dashed lines) (**C**); SMRT2 binding to LXRβ (solid lines) (**D**). Data shown mean and SD from at least triplicate measurements. (See **Fig. S6** for alternative layout).

A subset of ligands elicited strong agonist responses (high potency and efficacy for corepressor displacement) that were not strongly dependent on LXR isoform nor the identity of the corepressor: T0, RGX-104, GSK3987, and perhaps surprisingly, SR9238 that often displays the properties of an inverse agonist. AZ876 showed similar activity, with selectivity for corepressor displacement from LXRα (**Fig. 7**), the same isoform selectivity seen for coactivator recruitment (**Fig. 6**). MK9 and ALX101 elicited similar responses to the “strong agonists” in displacement of corepressors from LXRβ; however, these ligands were less effective or ineffective in displacement of corepressors from LXRα. BE1218 was less efficacious in general and was ineffective in corepressor displacement from LXRα. Thus, several LXR ligands only stabilize conformations of LXRα that allow corepressor binding.

A strong agonist is expected to potently destabilize corepressor binding to LXR. In this context, SR9238 has strong agonist properties (**Fig. S6Q**), which is perhaps surprising, since it is reported to act as an inverse agonist. In contrast to corepressor displacement, SR9238 was unable to recruit coactivators to LXRα and very weakly supported coactivator recruitment to LXRβ. Thus, SR9238 displays unique interactions with LXR, strongly displacing corepressors and with little effect on coactivator recruitment. SR9238 cannot stabilize a conformation of LXR that allows corepressor binding, nor one that stabilizes coactivator binding. The conformation stabilized by SR9238 also cannot be that of the apo-receptor, because the unliganded LXR apo-receptor binds corepressors.

Based on CRT analysis of coregulator binding, SR9238 would be predicted to act as an antagonist; therefore, it is useful to compare with GSK2033, universally described as an antagonist. GSK2033 stabilizes a conformation of LXR that is incompatible with coactivator binding and equally GSK2033 does not stabilize a conformation that disrupts corepressor binding to LXRα (**Fig. S6P**). GSK2033 does selectively displace SMRT2 from LXRβ. Two interesting conclusions result from this CRT analysis: 1) GSK2033 is selectively able to displace corepressors from LXRβ; and 2) GSK2033 is expected to act as a pure antagonist of LXRα-mediated transcription. In contrast, strong corepressor destabilization by SR9238 resulting in de-repression of LXR, is expected to activate LXR-mediated transcription. CRT data clearly differentiates SR9238 from the antagonist GSK2033.

Several ligands were unable to displace corepressors from LXR; and XL041 stabilized LXR complexes with corepressor bound, as shown by the concentration-dependent increase in the TR-FRET signal (**Fig. S6K**). Similar behavior was seen for the structurally related XL652 in stabilization of SMRT2 corepressor binding to LXRα; although, XL652 did not perturb SMRT2 binding to LXRβ and weakly destabilized the NCOR2 complex with both LXR isoforms (**Fig. S6L**). XL041 binding induced LXRα conformations that strongly stabilized the corepressor-bound complexes. The signal associated with corepressor binding to LXRβ was not increased to the same degree; furthermore, the signal associated with SMRT2 binding was increased to a higher maximal response than that for NCOR2 binding.

In summary, corepressor binding to LXR gave a much more diverse set of ligand-dependent responses in CRT, compared to those observed for coactivator binding; primarily, because unliganded LXR binds corepressors but not coactivators. Of the observations that stand out from the CRT corepressor binding analysis, the diverse response from the biphenyl sulfone ligands is perhaps the most surprising, with a spectrum of corepressor binding/stabilization to displacement/destabilization: XL041 > XL652 > GSK2033 > BE1218 > SR9238.

Simplistically, if expulsion of the corepressor from the LXRα complex is correlated with hepatic LXR activation and lipogenesis, AZ876, GSK3987, T0, RGX-104, and SR9238 would be predicted to be lipogenic. Derepression cannot be the only lipogenic mechanism in HepG2 cells, because SR9238 acts as an inverse agonist at < 3 µM concentration. Therefore, corepressor binding alone cannot be considered in the absence of coactivator recruitment.

### Synchronous coregulator recruitment measured by precision CRT

We established a precision CRT (pCRT) approach combining classical monoplexed CRT with a multiplexed, one donor/two-acceptor system allowing for simultaneous measurement of corepressor dissociation and coactivator recruitment. Terbium has multiple emission wavelengths therefore it is an ideal fluorescence donor. Utilizing two spectrally different acceptor fluorophores such as XL665 and fluorescein allows measurement of two signals in the same assay as the fluorophore acceptor emissions do not overlap. The prototype pCRT methodology utilized as corepressor, NCoR2-fluorescein, and as coactivator, the same construct, biotin-SRC1, used to determine LXR isoform selectivity (**Fig. 2**). The pCRT assay was optimized for a total volume of 12 µL incubating LXR (4 nM), anti-GST-Tb (0.4 nM), SA-XL665 (9.44 nM), SRC1-btn (200 nM) and NCoR2-FL (200 nM), measuring TR-FRET emission ratios at 520/490 nm and 665/615 nm. Coregulators were in excess and concentrations of assay components were kept constant while ligand concentration-response was examined.

It is clear from the pCRT curves that the varied response of corepressor NCoR2 to ligand binding seen in monoplexed CRT (**Fig. 7**) is lost in the presence of coactivator SRC1 (**Fig. 8**). For every ligand studied by pCRT, ligand binding induced a conformation of LXR compatible with coactivator recruitment and concomitant corepressor displacement. The synchronous exchange of corepressor by coactivator leads to pCRT binding curves in which the potency for ligand-induced binding of SRC1 is almost identical to that for ligand-induced displacement of NCOR2. For example: (1) LXR-623 does not displace NCOR2 from LXRα in the absence of SRC1; (2) DMHCA does not displace NCOR2 from LXRβ in the absence of SRC1; and (3) although GW3965 does displace NCOR2 from both LXR isoforms, the concentration-response for GW3965 shows a left-shift in the NCOR2 displacement curve of 100-fold (**Fig. 8**). In contrast to NCOR2 response, the SRC1 binding curves show only small shifts in in the presence of NCOR2 from pCRT analysis (**Fig. 8**). For LXR, the correlation between potency for SRC1 recruitment in monoplex and multiplexed CRT is excellent (**Fig. 4M-N**).

**Fig. 8.**
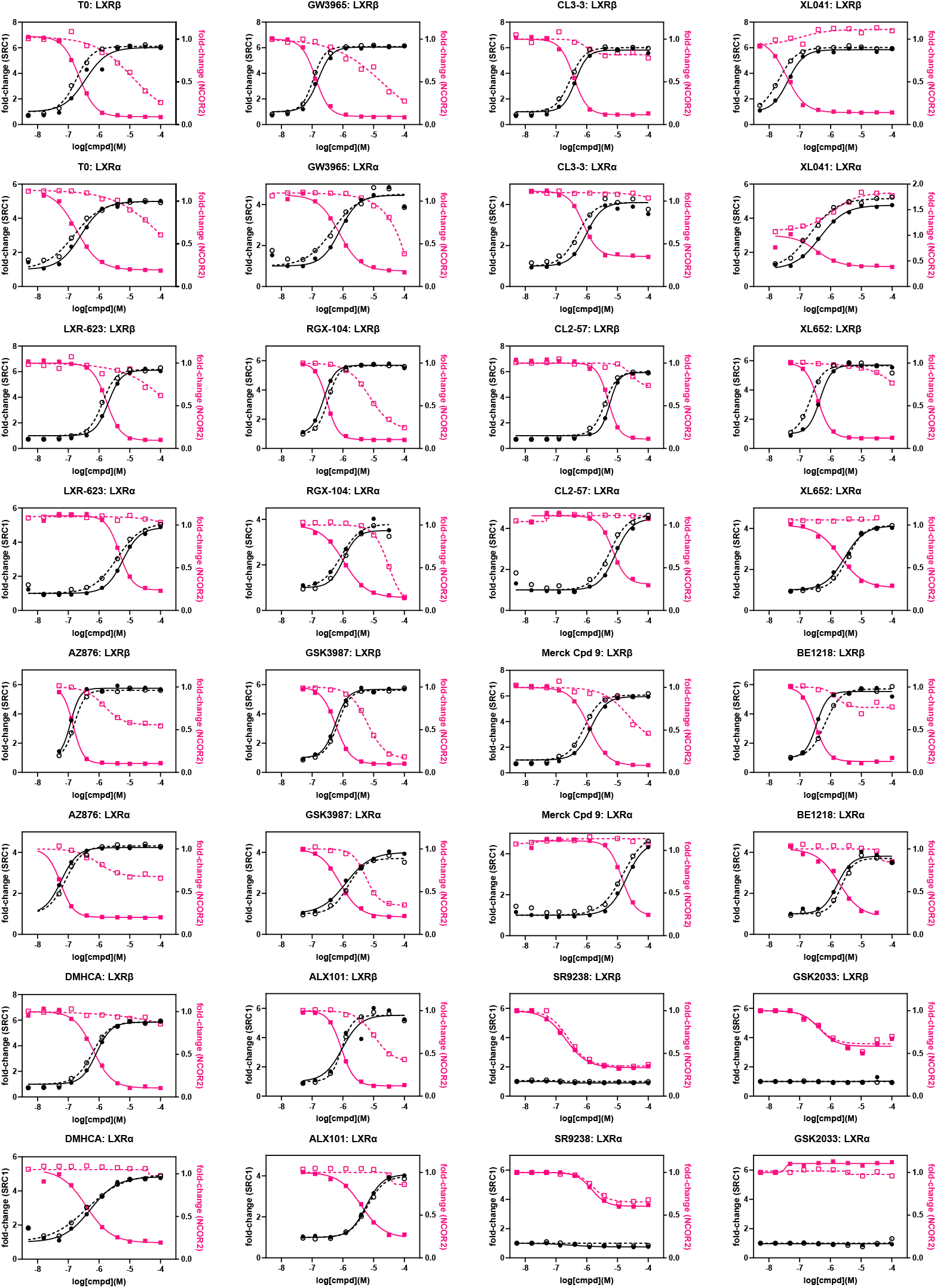
Ligand-induced conformational change to LXR causing corepressor (NCOR2) replacement by coactivator (SRC1) measured by pCRT. Concentration-response from CRT for SRC1 binding relative to apo-LXR (black dashed line) and NCOR2 binding relative to apo-LXR (magenta dashed line). Concentration-response from pCRT for SRC1 binding relative to apo-LXR (black solid line) and NCOR2 binding relative to apo-LXR (magenta solid line) with titration of ligand into solution of LXR, NCOR2, and SRC1. Data show mean of triplicate measurements.

Titration of the biphenyl sulfone ligands, XL041 and XL652, in the absence of SRC1 induced binding of NCOR2 to LXRα (for both ligands) and to LXRβ (XL041 only) (**Fig. 7**). From pCRT analysis, the presence of SRC1 causes ligand-induced displacement of NCOR2 from LXR. For LXRβ, in the presence of SRC1, the corepressor is fully displaced; whereas, the maximal signal for NCOR2-bound LXRα, is significantly higher than zero in the presence of saturating XL041 (**Fig. 8**) indicative of an equilibrium for the repressed XL041:NCOR2:LXRα and activated XL041:NCOR2:LXRα complexes.

CRT analysis of coactivator binding has often been used to determine ligand potency, efficacy, and isoform selectivity for LXR. CRT analysis of corepressors is more rarely studied. Concentration-dependence of four LXR ligands (T0, GW3965, LN6500 and 22(*R*)-hydroxy-cholesterol) was studied by CRT for coactivator NCoA3 and separately for NCOR1.^52^ Weak recruitment of NCOR1 to LXRα was reported for GW3965 (> 1 µM). This study was supported by cellular two-hybrid reporter assays in HepG2 and THP1 cells. T0 and GW3965 were classed as full and partial agonists, respectively. In our study, GW3965 and its methylated derivative, RGX-104, behaved as full agonists in CRT, with the exception of coactivator (PGC1α, SRC2, D22, TRAP220) recruitment to LXRα (**Figs 6, S6**). Both agonists displaced NCOR2 and SMRT2 from LXR. In contrast, two biphenyl sulfone LXR agonists (XL041, XL265) strongly stabilized the corepressor-bound conformation.

### Correlation of CRT signatures with phenotype

With the knowledge that, in pCRT, corepressors bind to the aporeceptor; whereas coactivator binding requires a ligand-induced conformational change, we can categorize all ligands using their CRT profiles. Ligands induce 3 LXR conformational ensembles with CoR (destabilizing D, stabilizing S, neutral N) and only 2 conformations with CoA (stabilizing S, neutral N) (**Table 1**). In categorizing ligands based upon these CRT characteristics, labels are needed.

**Table 1.** Coregulator stabilization signatures.

| Table 1. Coregulator stabilization signatures |  |  |  |  |
| --- | --- | --- | --- | --- |
| CRT | CoA | CoR | LXRα Derepression | LXRβ Canonical |
| Full agonist | S | D | Transcription | Transcription |
| Weak Agonist | S | N | Transcription | Transcription |
| Partial agonist | S | S | Partial | Transcription |
| Reverse Agonist | N | D | Transcription | Antagonist |
| Hard Antagonist | N | N | None | Antagonist |
| Soft Antagonist | N | S | None | Antagonist |
| S stabilized; D destabilized; N neutral |  |  |  |  |

The nomenclature for NR ligands often lacks precision and differs across NR classes. SERM (a subset of selective NR modulator) is used to describe varied families of ER ligands that show tissue-selective agonist and/or antagonist actions. Unfortunately, “partial agonist” is also widely used to describe SERMs, even though its use is usually pharmacologically problematic and biased agonist may be a more accurate label.^124^ The majority of reported ER ligands are SERMs, even some that cause ER degradation, because they are transcriptionally active. Consequently, the term “pure antagonist” (PA) has been used to differentiate transcriptionally null ligands^125^; although, pure antagonist/antiestrogen was originally introduced to describe antagonism of both AF1 and AF2 functions.^90^

Elegant work by Griffin’s team on RAR-related orphan receptor C (RORɣ) is interesting, because it used a combination of HDX-MS and CRT and labelled categories of RORɣ ligands.^126^ In addition to full agonist, “silent agonist” was introduced to include endogenous and synthetic partial agonists; although, by definition, partial agonists should antagonize full agonists. On the antagonist side of the spectrum, “active antagonist” was used to describe ligands that reduce cellular activity to baseline; and “inverse agonist” for ligands that reduce cellular transcription below baseline and induce recruitment of corepressors. Curiously, inverse agonist has almost never been used to describe ER ligands and is used frequently for other NR classes, mostly for ligands that reduce transcription below baseline, without any evidence for corepressor recruitment. GSK2033 and SR9238 show inverse agonist activity in cells (**Figs 3, 5**); however, neither is capable of recruiting SMRT2 or NCOR2 to LXR (**Fig. 7**).

We selected labels that were specific for the ligand signatures derived from pCRT data, mindful of the myriad and alternative uses of partial and inverse agonist: full agonist (FA), weak agonist (WA), partial agonist (PA), reverse agonist (RA), soft antagonist (SA), and hard antagonist (HA). The category for each ligand is not always identical for LXRα vs. LXRβ. Similarly, the phenotype associated with each classification is isoform dependent. Recruitment of CoA to LXRβ by FA, WA, and PA will activate transcription; whereas, lack of recruitment by SA, HA, and RA will antagonize transcription (**Table 1**). Stabilization of CoA and/or destabilization of CoR bound to LXRα is a derepression mechanism leading to activation/transcription for FA, WA, and RA; whereas partial activation is expected for PA.

To determine which LXR ligands fit with these theoretical CRT signatures, concentration-response curves were interpolated to measure normalized response at 1 µM ligand concentration and plotted as a heat map (**Fig. 9A**). Selecting response at 1 µM ligand is pharmacologically relevant and takes account of potency and efficacy. As benchmark lig- and, the response to T0 is uniform, without LXR isoform selectivity and with strong induction of ABCA1 and activation of SRE in cell cultures. Across the other 13 ligands with agonist activity, all are LXRβ-selective, with the exception of AZ876 that is LXRα-selective (leading to differentiation of AZ876 in hierarchical clustering analysis (**Fig. 4I**).

**Fig. 9.**
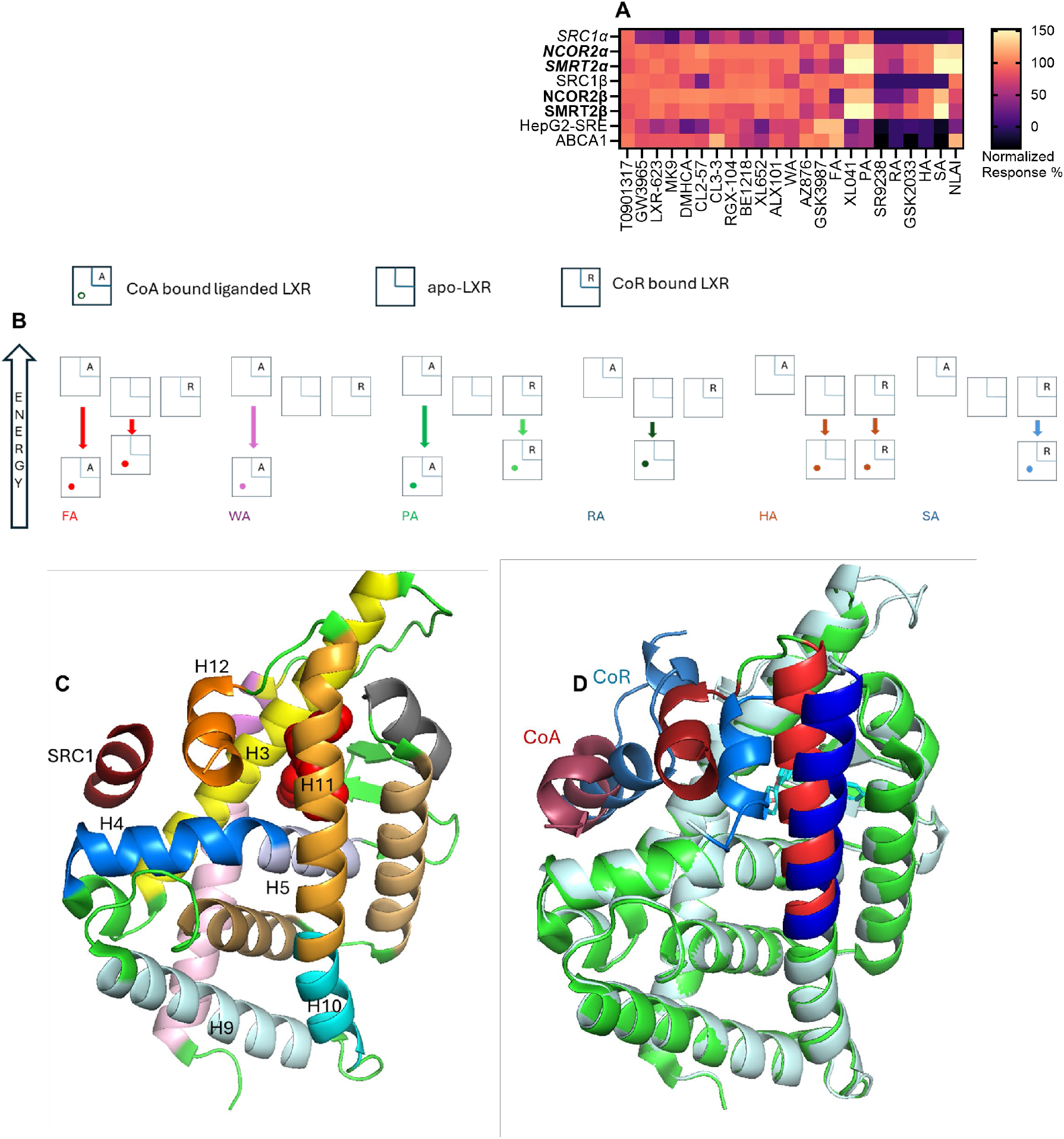
CRT signatures and correlation with stabilization and structure of LXR:coregulator complexes. (Table 1) Categorization of LXR ligands as agonists and antagonists conforming to CRT and pCRT signatures. **(A)** Heat map of responses (ligand concentration at 1 µM) in cell-free and cell-based assays to LXR ligands together with theoretical response to agonist and antagonist ligand signatures. **(B)** Relative energy perturbations of apo and coregulator-bound LXR caused by each class of agonist and antagonist identified and classified in Table 1. **(C)** Co-crystal structure of LXRα:SRC1-2 co-crystal with GW3965 (PDB 3IPQ) showing key helices. **(D)** Superposition of LXRα:SRC1-2:GW3965 co-srystal with structure of LXRα:SMRT2 simulated using Alphafold, showing the significant structural perturbation of H11 and H12 on ligand binding (red shades) compared to the ligand-free LXRα (blue shades).

The pCRT signatures of the three agonist and three antagonist classifications were modelled in the same heat map using signatures defined in **Table 1** and the measured responses to the real LXR ligands in our study (**Table S3**). There are several ligands with weak agonist (WA) signatures and GSK3987 fits a full agonist (FA) profile. XL041 has the signature of a partial agonist (PA), which matches its cellular phenotype. GSK2033 and SR9238 match the CRT signatures of a hard antagonist (HA) and inverse agonist (RA), respectively. We hypothesized that the signature of an NLAI, which by definition induces ABCA1 but does not activate SRE, should reflect a CRT profile of LXRβ activation without activating LXRα: this leads to a theoretical NLAI profile, shown in **Fig. 9A**, which is not an exact match for any theoretical or experimental LXR ligand.

To give these signatures and labels a more thermodynamic basis, we can take learnings from CRT/pCRT measurements: (i) CoA-bound LXR is of relatively high energy, since it is not observed in absence of ligand; (ii) apo-LXR exists in equilibrium with CoR bound LXR, since CoR-bound LXR is observed in absence of ligand; (iii) The ligand-bound CoA:LXR complex is lowest energy, because CoA binding leads to full or partial CoR displacement. The relative stabilization and relative energy associated with each complex therefore follows (**Fig. 9B**). Thus, a reverse agonist (RA) CRT signature requires the RA to stabilize apo-LXR and a partial agonist (PA) stabilizes both coregulator-bound complexes.

### Implications for discovery of nonlipogenic LXR ligands

As outlined in the Introduction, Belorusova et al., carried out multicomponent analysis on an in-house chemical series of 9 LXR ligands, T0, GW3965, LXR-623, XL041, and AZ876, claiming identification of an “escape route from the adverse lipogenic effects of LXR ligands”. ^50^ Although the study was limited by binary definitions of *in vivo* lipogenicity, the in-depth HDX-MS analysis led to inferences on ligand-induced conformational change that might be associated with lipogenesis. Not unexpectedly, given the complexity of the problem, simple principles for NLAI design were not revealed. It was concluded that selection of ligands with “modest potencies” in cell-based transcriptional assays would lead to reduced lipogenesis.

Correlation analysis driven by HDX-MS data inferred the importance of helix-12 (H12) in activating a lipogenic transcriptional program. H12 is the key structural element of Activation Function-2 (AF-2), often described as a ligand-dependent switch and providing a binding surface for coactivator recruitment in the agonist-induced conformation (**Fig. 9C**). There are no published crystal structures of corepressor-bound LXRα, with most PDB entries being liganded coactivator-bound LXRβ. To emphasize the conformational change in LXRα induced by agonist binding, we modelled SMRT2-bound apo-LXRα using Alphafold and superimposed the corepressed apo-structure with a SRC1-bound agonist co-crystal structure (PDB 3IPQ **Fig. 9D**). This illustration shows the CoR, in the absence of ligand, bound to a surface that in the presence of ligand is occupied by H12. A ligand-induced conformation that stabilizes CoR binding to LXR cannot result from direct displacement of H12 by ligand.

Helix-3 (H3) forms part of the coactivator binding surface and the ligand-binding pocket. H3 contributes residues that interact directly with ligands. Helix 4 (H4) interacts with H3 and other helices to form the hydrophobic core, stabilize coactivator-bound conformations, and indirectly to influence receptor dimerization through helix-9 (H9). In LXR, helix-5 (H5) interacts directly with ligands and supports a more flexible ligand-binding pocket than in many other NRs. HDX-MS analysis of liganded LXRα structures correlated stabilization of H3 and H5 with selective *ABCA1* induction. It was speculated that ligand interactions with H3 and H5 could induce a conformation leading to corepressor dissociation, which could lead to selectivity owing to the greater flexibility of H3 in LXRα. ^50^ Conversely, ligands that stabilized H12, itself, were correlated with lipogenesis. Similarly, direct stabilization of H12 of PPARγ is associated with full agonism; whereas, partial agonists do not directly stabilize H12, which leads to biased agonism.^124^

As in our study, Belorusova et al noted that the properties of XL014, were differentiated from other LXR ligands. The HDX-MS analysis showed XL014 to act as an antagonist: i.e. XL014 did not induce an agonist conformation of H12. XL041 is generally described as an LXRβ-selective partial agonist.^127^ Our CRT analysis of coactivator binding (**Figs 6, S5**) reflects LXRβ-selective partial agonist activity, with the exception of SRC1 CRT data. In cell-based phenotypic assays, XL041 is a potent partial agonist with selectivity (at concentration ≤ 1 µM) for induction of ABCA1 over SRE activation (**Figs 3, 5**). Taken together with the XL041-induced stabilization of corepressor-bound LXR (**Fig. 7**) the overall CRT signature of XL041 uniquely predicts partial agonist activity (**Fig. 9**).

XL041 stabilizes corepressor binding to LXRα (**Fig. 7**); although, in the presence of coactivator, SRC1, the corepressor is dissociated (**Fig. 8**). In our study, the biphenyl sulfone class of ligands yielded diverse responses in cell-free and cell-based assays, suggesting that this chemotype can be manipulated to enhance isoform and coregulator selectivity. The CRT profile of XL652, a close homologue of XL041, includes corepressor stabilization for LXRα (**Fig. 7**) and a response in HepG2 cells that shows little or no activation of SRE at submicromolar concentrations and more than 100-fold weaker potency for ABCA1 induction compared to XL041 (**Figs 3, 5**). The phenotype of ALX101 is a full agonist towards ABCA1 induction and a partial agonist of SRE activation; whereas, BE1218 has a partial agonist phenotype in both cell-based assays. The partial agonist phenotype is compatible with a relative inability to destabilize the corepressor-bound LXRα complex (**Fig. S6**).

GSK2033 and SR9238 are very close structural congeners of BE1218, yet the response in phenotypic assays is quite different, with BE1218 a partial agonist, SR9238 displaying an inverse agonist and GSK2033 a mixed antagonist/inverse agonist phenotype (**Figs 3**,**5**). In CRT, both GSK2033 and SR9238 displace SRC1 and 24HC from an activated LXRβ complex (**Fig. 2**) but GSK2033 more closely resembles BE1218 in its weak effects on corepressor-bound LXRα (**Fig. S6**). There is scope for further exploration of this chemotype using CRT/pCRT with a focus on interactions with the LXRα corepressor complex.

### Limitations of the present study

Measurement of absolute affinity for coregulator binding to LXR by TR-FRET or other methods is complementary to the CRT approach used in this study, which measures relative potency and efficacy for ligand-induced coregulator binding. Absolute measurements require saturating ligand concentrations, which is problematic for weak LXR ligands, which by their nature are hydrophobic with limited aqueous solubility. Elegant studies measuring PPAR/coregulator binding free energy by fluorescence polarization have incorporated RXR heterodimers, full length NRs, and oligonucleotides.^91, 124^ These studies demonstrate that key aspects of ligand-induced coregulator binding are replicated by the reductionist system of NR-LBD with short coregulator peptides as used in the present work.

## Conclusions

Partial agonists are often described as a desirable therapeutic modality with anticipated efficacy and improved safety; although, the rationale for this improved therapeutic window is often intuitive rather than quantitative. The traditional medicinal chemistry approach to optimization of small molecule therapeutics is to increase potency and improve selectivity against an off-target: the approach in LXR agonist design has invariably been LXRβ versus LXRα isoform selectivity. A nominated drug candidate will also maximize target exposure. XL041 ticks all the boxes as a potent, LXRβ-selective ligand with partial agonist response in many *in vitro* assays. In human volunteers, XL041 caused a non-linear, dose-dependent increase in blood ABCA1 and plasma TG and LDL-C, with maximal response reached at approximately 2.5 mg/kg p.o. At this dose, the drug concentration in plasma on Day 14 was C_max_ ≈ 30 nM. Levels of TG and LDL-C returned to baseline after drug withdrawal; however, with T½ ≈ 13 hours and the high potency of XL041, target exposure would be expected to be persistent. Limiting target exposure and potency may be required to attenuate activation of lipogenesis even by partial agonists. The clinical observations on the biphenyl sulfone XL041 are compatible with *in vitro* data reported by others and in this present study.

Preclinical animal studies have shown that enhancing ABCA1 activity with an LXR agonist could be an effective therapeutic strategy in a wide range of human conditions, including diabetes, cardiovascular disease, kidney failure, retinal degeneration, and neurodegenerative diseases, including ADRD.^38, 128-130^ In preclinical proof-of-concept studies T0, RGX104 and GW3965 continue to be used; for example, GW3965 in a tauopathy model of pathologic glial cell activation associated with AD.^69^ We have reported beneficial effects of CL3-3 in FAD mice expressing human *APOE4*, the major genetic risk factor for AD.^131^ We observe a strong correlation between cell-free stabilization of CoA:LXRβ complexes versus induction of ABCA1 in cells, supportive of the therapeutic focus on LXRβ selectivity. Hepatic lipogenesis as a side effect, caused by LXR activation in the liver, cannot be tolerated in many chronic diseases. Hepatic lipogenesis, observed in the XL041 clinical trial, is not tolerable in a population prone to atherosclerosis, the disease target of XL041. The diverse responses of LXR ligands reflected in CoR:LXRα stabilization and SRE activation in HepG2 cells suggests that fine tuning of ligands with respect to LXRα binding is a route to attenuation of unwanted lipogenesis.

## Experimental Section

### Cell Culture

CCF-STTG1 and HepG2 cell lines were purchased from ATCC and cultured per the supplier’s instructions. All cultures were maintained at 37°C in a humidified environment containing 5% CO_2_. CCF-STTG1 cultures were grown in RPMI-1640 media (ATCC) and HepG2 in Eagles Modified Essential Medium (EMEM, ATCC 30-2003) supplemented with 10% fetal bovine serum (FBS, Gibco A52567-01) and 1% Penicillin/Streptomycin (P/S, Gibco 15140-122).

### Luciferase reporter assays

#### ABCA1-luc assay

Low passage CCF-STTG1 cells (20,000 cells/well) were seeded in 96-well plates in media containing charcoal-stripped FBS and incubated at 37 °C and 5% CO_2_ for 24 hours prior to a 24-hour treatment. All compounds were dissolved in DMSO, and final DMSO concentrations never exceeded 1%. After a 24-hour incubation period, add a volume of ONE-GLO™EX Reagent that is equal to the volume of culture medium in each well and incubate for 3 to 5 minutes on the shaker. The luminescent activity of compounds (X to Y µM, 10-fold dilution) was examined using the ONE-GLO™EX assay (Promega). The data was normalized to DMSO. The assays were performed in duplicates for each concentration, and the EC50 values were determined from non-linear regression analysis of the dose-response curve generated in GraphPad Prism 9.

#### SRE-luc assay

HepG2 cells were cotransfected with SRE (LDLR)-NlucP SuperPiggyBac Transposon and a separate Transposase plasmid (System Biosciences). Cells were expanded for multiple passages followed by selection with puromycin (1ug/ml). Stable cells constitutively express GFP allowed for FACS enrichment and/or cell cloning. Initially, stable pools were compared to low expressing and high expressing enriched pools. The optimal pool was selected based on assay performance (Z’) and was grown and maintained as above, with the addition of 1.0 μg/mL of puromycin biweekly.

Cells were split into flasks containing CS medium (EMEM supplemented with 10% charcoal stripped FBS (Gibco A52567-01) and 1% P/S and grown at 37°C /5% CO_2_ for 48 hours before plating. Cells were plated in 384-well, white clear tissue culture plates (Greiner 781098) and allowed to grow at 37°C /5% CO_2_ for 24 hours before medium was aspirated and compound dilutions prepared in CS medium were added. Following a 24-hour incubation, the medium was aspirated, and cells were lysed/luminescence quantified using the Nano-Glo Luciferase Assay System (Promega N1130). Luminescence was measured on a CLARIOstar® Plus multi-mode plate reader (BMG Labtech, Ortenberg, Germany). Experimental data were normalized to controls and fit with a four parameter Hill equation using Graphpad Prism software (Boston, MA) to determine EC_50_ values.

#### qPCR

HepG2 cells were seeded in 12-well plates and treated with compounds for 24 hours. Following treatment, cells were washed once with PBS and then lysed in 350 μL of RLT buffer (Qiagen). Lysates were immediately frozen at −80 °C until RNA extraction. Total RNA was extracted using RNeasy Plus Mini columns (Qiagen) according to the manufacturer’s instructions. RNA concentration and purity were assessed using the A260/A280 ratio measured on a Synergy Neo2 plate reader (BioTek). For cDNA synthesis, 2 μg of total RNA was reverse transcribed using SuperScript III reverse transcriptase (Invitrogen), oligo(dT)12–18 primers, and dNTP mix, following the manufacturer’s protocol. Quantitative PCR was performed using TaqMan Gene Expression Master Mix (Invitrogen) with 2 μL of cDNA and gene-specific TaqMan probes in a 20 μL reaction volume per well. The following TaqMan assays were used: ABCA1: Hs01059101_m1, SREBF1: Hs01088691_m1, SCD: Hs01682761_m1, FASN: Hs01005622_m1, and HPRT1 (endogenous control): Hs02800695_m1. qPCR was performed on a StepOne Real-Time PCR System (Applied Biosystems), and gene expression was quantified using the ΔΔCt method.

### Coactivator recruitment TR-FRET/HTRF (CRT)

#### LXRα/β

To determine compound EC50 values, 10mM compound stocks were prepared in DMSO. An initial 8-concentration, 3-fold dilution series in DMSO was performed in 96-well v-bottom, polypropylene plates at 100x the final desired concentration. Intermediate dilution plates were prepared by diluting the 100x DMSO stocks 25-fold in assay buffer (50mM Tris, pH 7.4, 50mM KCl, 0.05% Triton-X 100, 10% glycerol, 0.1% BSA-(added fresh), 5mM DTT (added fresh)), creating a 4x stock. The following assay components were prepared individually at a 4x final concentration in assay buffer: (1) LXRα/β - GST tag (Thermo Fisher Scientific PV4657/PV PV4660); (2) a mixture of anti-GST-Tb (CisBio/Perkin Elmer 61GSTTLA, lot 14A) and SA-XL665 (CisBio/Perkin Elmer 610SAXLB); (3) SRC1-btn (AnaSpec AS62152/Fisher Scientific NC1565694). A 3uL aliquot of the 4x ligand was then added to each well of the 384-well small volume non-binding black assay plate (Greiner 784900) followed by a 3uL aliquot of each reaction component for a total volume of 12uL. Control wells contained all components minus ligands, replaced by positive control T0 (Enzo 270-309-M010/VWR 89147-616) or DMSO. Incubated at room temperature for 1hr before measurement of TR-FRET. Final concentrations of assay components LXR *α*/*β* (4nM), anti-GST-Tb (0.4nM), SA-XL665(9.44nM), and SRC1-btn (100nM). TR-FRET measurements were performed on a CLARIOstar® Plus multi-mode plate reader (BMG Labtech, Ortenberg, Germany), excitation 337nm (laser), emission1 615nm and emission2 665nm. The signal is expressed as a ratio of TR-FRET measurements at 665 nm/615nm × 1000. Experimental data were normalized to controls and fit with a four parameter Hill equation using Graphpad Prism software (Boston, MA) to determine EC50s. Full details for other NR CRT assays are included in **Supporting Information**.

#### LXR (coregulator selectivity)

For all assays, ligands were first prepared at 100x of desired final concentration in DMSO. Ligands were serially diluted in a 3x manner in 96-well polypropylene plate. The ligands were then diluted 4x final concentration in assay buffer. The following assay components were prepared individually at a 4x final concentration in assay buffer: (1) LXRα/β -GST tag (Thermo Fisher Scientific PV4657/PV PV4660); (2) a mixture of anti-GST-Tb (Cis-Bio/Perkin Elmer 61GSTTLA, lot 14A) (3) coregulator-FL. A 3uL aliquot of the 4x ligand was then added to each well of the 384-well small volume non-binding black assay plate (Greiner 784900) followed by a 3uL aliquot of each reaction component for a total volume of 12uL. Control wells contained all components minus ligands, replaced by positive control T0 (Enzo 270-309-M010/VWR 89147-616) or DMSO. Incubated at room temperature for 1hr before measurement of TR-FRET. Final concentrations of assay components LXR*α*/*β* (3nM), anti-GST-Tb (0.4nM), and coregulator-FL (200nM). TR-FRET measurements were performed on a CLARIOstar® Plus multi-mode plate reader (BMG Labtech, Ortenberg, Germany), excitation 337 nm (laser), emission1 520nm and emission2 490 nm. The signal is expressed as a ratio of TR-FRET measurements at 520nm/490nm × 1000. Experimental data were normalized to controls and fit with a four parameter Hill equation using Graphpad Prism software (Boston, MA) to determine EC50s.

#### Multiplex LXR pCRT one donor/two-acceptor fluorophore

For all assays, ligands were first prepared at 100x of desired final concentration in DMSO. Ligands were serially diluted in a 3x manner in 96-well polypropylene plates. The ligands were then diluted to a 4x final concentration in assay buffer. The following assay components were prepared individually at a 4x final concentration in assay buffer: (1) LXR*α*/*β*; (2) a mixture of anti-GST-Tb and SA-XL665; (3) a mixture of SRC1-btn and NCoR2-FL. A 3uL aliquot of the 4x ligand was then added to each well of the 384-well black assay plate followed by a 3uL aliquot of each reaction component for a total volume of 12uL. Incubated at room temperature for 1hr before measurement of TR-FRET. Final concentrations of assay components were LXR*α*/*β* (4 nM), anti-GST-Tb (0.4 nM), SA-XL665(9.44 nM), SRC1-btn (200 nM) and NCoR2-FL (200 nM). TR-FRET measurements were performed on a CLARIOstar® Plus multi-mode plate reader (BMG Labtech, Ortenberg, Germany). Two different measurements were taken. The first measurement was excitation 337nm (laser), emission1 520nm and emission2 490nm. The signal is expressed as a ratio of TR-FRET measurements at 520 nm/490nm × 1000. The second measurement was excitation 337nm (laser), emission1 615nm and emission2 665nm. The signal is expressed as a ratio of TR-FRET measurements at 665 nm/615nm × 1000. Experimental data were normalized to controls and fit with a four parameter Hill equation using Graphpad Prism software (Boston, MA) to determine EC_50_ values.

#### Modelling. Structural Alignment and Model Generation Using PyMOL and AlphaFold

The SMRT2 peptide sequence (CPSSHSSLTERHKILHRLLQEGSPS) and FASTA sequence of LXRα (PDB 3IPQ) were submitted together to the AlphaFold server using the “Sequences” tool with the prediction mode set to “Protein” yo yield a predicted model of the LXRα–SMRT2 complex. To evaluate this model, we aligned it with the LXRα:SRC1-2 co-crystal with GW3965.

#### Chemical synthesis

Detailed methods are provided in Supporting Information for CL2-57, CL3-3 and Merck Compound 9, including characterization and purity.

## ASSOCIATED CONTENT

### Supporting Information

Tables S1 and S2 showing NR Panel assay components and CRT peptide sequences. Fig S1 showing NR Panel responses. Fig S2 showing control reproducibility in cellular experiments. Fig S3 showing additional correlation plots of cellular data and LXR isoform selectivity. Fig S4 showing full dose response curves for HepG2 data. Fig S5 showing coactivator correlation plots. Fig S6 showing all coactivator responses for ligands. Fig S7 corepressor responses separated by individual ligands. Fig S8 heat map showing coregulator efficacy across ligands. Experimental details for NR Panel experiments and chemical synthesis. NMR spectra for synthesized compounds.

## AUTHOR INFORMATION

### Corresponding Author

#### Gregory R J Thatcher

Department of Pharmacology & Toxicology, R Ken Coit College of Pharmacy, Department of Chemistry & Biochemistry, Colleges of Science and Medicine, University of Arizona, Tucson, Arizona 85721, USA; orcid.org/0000-00027757-1739;

### Author contributions

GRJT conceived the project. MSL designed and coordinated the experimental approach. In vitro studies were conducted by MSL, MAB, ST, FA, MR, SRM, AA, SK, MS, CP and NM. Chemistry was conducted by GRV. The manuscript was written by GRJT and MSL with input from all authors.

#### Notes

GRJT is an inventor on patents owned by the University of Illinois.

## ABBREVIATIONS

(AD): Alzheimer’s disease
(ADRD): Alzheimer’s disease and related dementia
(Aβ): amyloid-β
(APOA1): Apolipoprotein A1
(APOE): Apolipoprotein E
(ABC): ATP-binding cassette transporter
(ACC): acetylCoA carboxylase
(AF-2): activation function-2
(APP): amyloid precursor protein
(BBB): blood-brain barrier
(CAR): constitutive androstane receptor
(CETP): cholesteryl ester transfer protein
(CNS): central nervous system
(CRT): coregulator time-resolved fluorescence energy transfer
(pCRT): precision CRT
(CSF): cerebrospinal fluid
(CYP): cytochrome p450
(CVD): cardiovascular disease
(DDIs): drugdrug interactions
(DMHCA): N.N-dimethyl-3*β*-hydroxycholenamide
(EMEM): eagles modified essential medium
(ER): estrogen receptor
(FA): full agonist
(FAS): fatty-acid synthase
(FAD): familial AD
(FXR): farnesoid X receptor
(FBS): fetal bovine serum
(HA): hard antagonist
(H3): helix-3
(H5): Helix-5
(H): helix-9
(H12): helix-12
(HDL): high-density lipoprotein
(HFD): high-fat diet
(HTS): highthroughput screen
(HDX-MS): H/D-exchange mass spectrometry
(IA): inverse agonist
(KO): knockout
(LDL): low-density lipoprotein
(LXR): liver X receptor
(MK9): Merck Compound 9
(NASH): non-alcoholic steatohepatitis
(NCOA1): nuclear receptor coactivator 1
(NCoR): nuclear receptor corepressor
(NHR): nuclear hormone receptor
(NLAI): nonlipogenic ABCA1 inducer
(OCA): obeticholic acid
(PA): partial agonist
(PPAR): peroxisome proliferator activated receptor
(PGC-1): PPAR*γ* coactivator 1
(PXR): pregnane X receptor
(P/S): penicillin/streptomycin
(RCT): reverse cholesterol transport
(RAR): retinoic acid receptor
(RXR): retinoid X receptor
(RA): reverse agonist
(SA): soft antagonist
(SERMs): selective estrogen receptor modulators
(SMRT): silencing mediator of retinoic acid and thryroid hormone receptor
(SRC-1): steroid receptor coactivator 1
(SRE): sterol response element
(SREBP1c): sterol-response element binding protein 1c
(SCAP): SREBP cleavage-activating protein
(SRE-luc): SRE promoter luciferase reporter assay
(TG): triglyceride
(TR-FRET): time-resolved fluorescence energy transfer
(TRAPs): thyroid hormone receptor-associated proteins
(T0): T0901317
(T2D): type-2 diabetes
(WA): weak agonist
(WT): wild-type
(VLDL-C): very low-density lipoprotein cholesterol
(24OHC): 24-hyrdoxycholesterol
(24HC): 24(*S*)hydroxycholesterol
(27OHC): 27-hydroxycholesterol

