## Supplemental Information for "In search of nonlipogenic ABCA1 inducers (NLAI): precision coregulator TR-FRET identifies diverse signatures for LXR ligands"

**Table S1. Summary of NR Panel Parameters.**

| Lanthascreen | | |
|---|---|---|
| **Receptor** | **Coactivator/Acceptor** | **Donor** |
| *FXR* | SRC2-2-fluorescein | $\alpha$-GST-Terbium |
| *PXR* | SRC1-4-fluorescein | $\alpha$-GST-Terbium |
| *PPAR$\delta$* | C33-fluorescein | $\alpha$-GST-Terbium |
| *RXR$\alpha$/$\beta$* | D22-fluorescein | $\alpha$-GST-Terbium |
| *RAR$\alpha$* | D22-fluorescein | $\alpha$-GST-Terbium |
| *RAR$\beta$* | SRC2-2-fluorescein | $\alpha$-GST-Terbium |

**Table S2. Coregulator Peptide Sequences**

| Vendor | Catalog No. | Coregulator | Sequence | MW (Da.) |
|---|---|---|---|---|
| Anaspec | AS-62152 | SRC1-btn | Biotin-CPSSHSSLTERHKILHRLLQEGSPS | 3026.6 |
| Thermo Fischer | PV4386 | FL-D22 | Fluorescein-LPYEGSLLLKLLRAPVEEV | 2499 |
| Thermo Fischer | PV4549 | FL-TRAP220/DRIP-1 | Fluorescein-NTKNHPMLMNLLKDNPAQD | 2554 |
| Thermo Fischer | PV4582 | FL-SRC1-4 | Fluorescein-GPQTPQAQQKSLLQQLLTE | 2465.9 |
| Thermo Fischer | PV4586 | FL-SRC2-2 | Fluorescein-LKEKHKILHRLLQDSSSPV | 2585 |
| Thermo Fischer | PV4421 | FL-PGC1a | Fluorescein-EAEEPSLLKKLLLAPANTQ | 2424 |

| | | | | |
|---|---|---|---|---|
| Thermo Fischer | PV4606 | FL-C33 | Fluorescein-HVEMHPLLMGLLMESQWGA | 2539 |
| Thermo Fischer | PV4624 | FL-NCoR ID2 | Fluorescein-DPASNLGLEDIIRKALMGSFDDK | 2865 |
| Thermo Fischer | PV4423 | FL-SMRT ID2 | Fluorescein -HASTNMGLEAIIRKALMGKYDQW | 2993 |

**Table S3. Data used in correlation plots and heatmaps.**

Table S3A: Potency (-log $EC_{50}$)

| | PGC | SRC2 | TRAP220 | D22 | PGC | SRC2 | TRAP220 | D22 | ABCA1 | SRE |
|---|---|---|---|---|---|---|---|---|---|---|
| T0901317 | 5.31034 | 5.18977 | 4.98758 | 5.71019 | 5.55533 | 5.53018 | 5.42447 | 5.63733 | 7.62342 | 7.63582 |
| GW3965 | 5.20419 | 5.14249 | 4.48825 | 4.79156 | 5.06717 | 5.25532 | 5.14655 | 5.15639 | 7.12901 | 6.36351 |
| XL041 | 5.57774 | 5.71422 | 5.77833 | 5.86646 | 7.06925 | 7.07779 | 7.06344 | 7.04735 | 9.18046 | 8.24718 |
| LXR-623 | 4.53835 | 4.40827 | 4.68952 | 4.85232 | 4.98005 | 4.87811 | 4.96417 | 5.03315 | 6.5236 | 5.8911 |
| Merck | 1.33809 | 2.68048 | 2.49431 | 3.94233 | 4.98005 | 4.84345 | 4.90448 | 5.07889 | 6.91686 | 6.64207 |
| DMHCA | | | | | | | | | 6.58687 | |
| CL2-57 | 4.80134 | 4.43344 | 5.33003 | 4.46395 | 5.13674 | 4.36191 | 5.38195 | 4.9165 | 6.09561 | 5.54485 |
| CL3-3 | 5.70885 | 5.62562 | 5.39729 | 5.66354 | 5.74642 | 5.79398 | 5.67572 | 5.78941 | 7.19586 | 6.84406 |
| RGX-104 | 5.5293 | 5.64936 | 5.39233 | 5.53091 | 5.85761 | 5.87844 | 5.69465 | 5.71851 | 7.19179 | 6.68867 |
| AZ876 | 6.8066 | 6.73049 | 6.49255 | 7.0271 | 6.44831 | 6.51856 | 6.17179 | 6.42262 | 7.96658 | 8.37345 |
| BE1218 | | | 5.44394 | 5.10325 | | | 5.99525 | 6.01601 | 8.04048 | 6.78516 |
| XL625 | 5.27344 | 4.85511 | 5.37397 | 5.47095 | 6.0765 | 6.17908 | 5.92775 | 6.02365 | 6.84741 | |
| ALX101 | 4.30751 | 0.19853 | 4.21954 | 4.73613 | 4.76751 | 4.72677 | 4.71783 | 5.34775 | 7.02411 | 5.99268 |
| GSK3987 | 5.49621 | 5.56687 | 5.96859 | 5.36917 | 5.66716 | 5.71806 | 6.15695 | 5.78331 | 7.34314 | 6.73166 |

Table S3B: Rel % activity at 1 uM used in Heat Map Analysis:

| | SRC1α | NCOR2α | SMRT2α | SRC1β | NCOR2β | SMRT2β | HepG2-SRE | ABCA1 |
|---|---|---|---|---|---|---|---|---|
| T0901317 | 93.37 | 91.70310528 | 83.05925029 | 97.84 | 92.3401757 | 84.98235327 | 91.33 | 99.73395993 |
| GW3965 | 37.86 | 99.05217728 | 92.36071088 | 95.64 | 92.2684161 | 92.51996642 | 71.74 | 81.39386888 |
| LXR-623 | 34.21107858 | 91.73073033 | 100 | 92.41087332 | 103.7005432 | 100.3928571 | 48.5261 | 83.67238173 |
| MK9 | 8.83084746 | 95.10137458 | 87.90804035 | 93.87254175 | 102.6843752 | 87.86368508 | 61.4664 | 83.9208158 |
| DMHCA | 59.95292422 | 100 | 99.37736369 | 75.40275393 | 106.3910195 | 98.67667443 | 18.8 | 85.48171348 |
| CL2-57 | 19.18666619 | 109.63873 | 99.98347371 | 27.25145715 | 107.4349509 | 99.35669185 | 34.43488083 | 80.44624312 |
| CL3-3 | 60.31092311 | 100 | 87.87649944 | 85.30087579 | 104.3122482 | 88.55637075 | 78.70930252 | 123.7078516 |
| RGX-104 | 71.25484176 | 99.78958921 | 93.44698343 | 100.1584679 | 103.279539 | 81.95589688 | 87.38555279 | 82.47792462 |
| BE1218 | 38.85784994 | 100 | 91.79031714 | 98.36273939 | 104.8578543 | 81.55595221 | 66.3 | 62.85320742 |
| XL652 | 59.04699685 | 100 | 97 | 100.793569 | 107.2985743 | 102.3253449 | 25 | 51.53916966 |
| ALX101 | 38.79156306 | 100 | 87.29628585 | 85.27073661 | 105.3830721 | 91.13807062 | 72.0265088 | 88.84231882 |
| WA | 70 | 100 | 100 | 70 | 100 | 100 | 70 | 70 |
| AZ876 | 101.3172196 | 79.44936072 | 46.06574835 | 98.95475658 | 78.35684164 | 63.48303759 | 87.95632601 | 106.7013798 |
| GSK3987 | 82.51218782 | 97.67143034 | 69.85630792 | 95.28317524 | 97.40247735 | 84.02254692 | 127.4639312 | 101.9248339 |
| FA | 101 | 64 | 46 | 100 | 22 | 62 | 127 | 123 |
| XL041 | 82.80769567 | 141.6 | 152.5 | 95.37246141 | 128.8722945 | 152.5495087 | 22.5 | 44.08501745 |
| PA | 101 | 142 | 152 | 100 | 128 | 152 | 50 | 50 |
| SR9238 | 0 | 64.52046255 | 64.47431302 | 0 | 21.90467677 | 69.21964091 | -22.158 | -34.0745463814932 |
| RA | 0 | 64 | 46 | 0 | 22 | 62 | 0 | 0 |
| GSK2033 | 0 | 97.67143034 | 92.90227612 | 0 | 57.69163298 | 93.36052342 | 5 | -33.2927421891987 |
| HA | 0 | 100 | 100 | 0 | 100 | 100 | 0 | 0 |
| SA | 0 | 142 | 152 | 0 | 128 | 152 | -22 | -34 |
| NLAI | 8.8 | 140 | 152 | 100 | 78 | 63 | 30 | 123 |

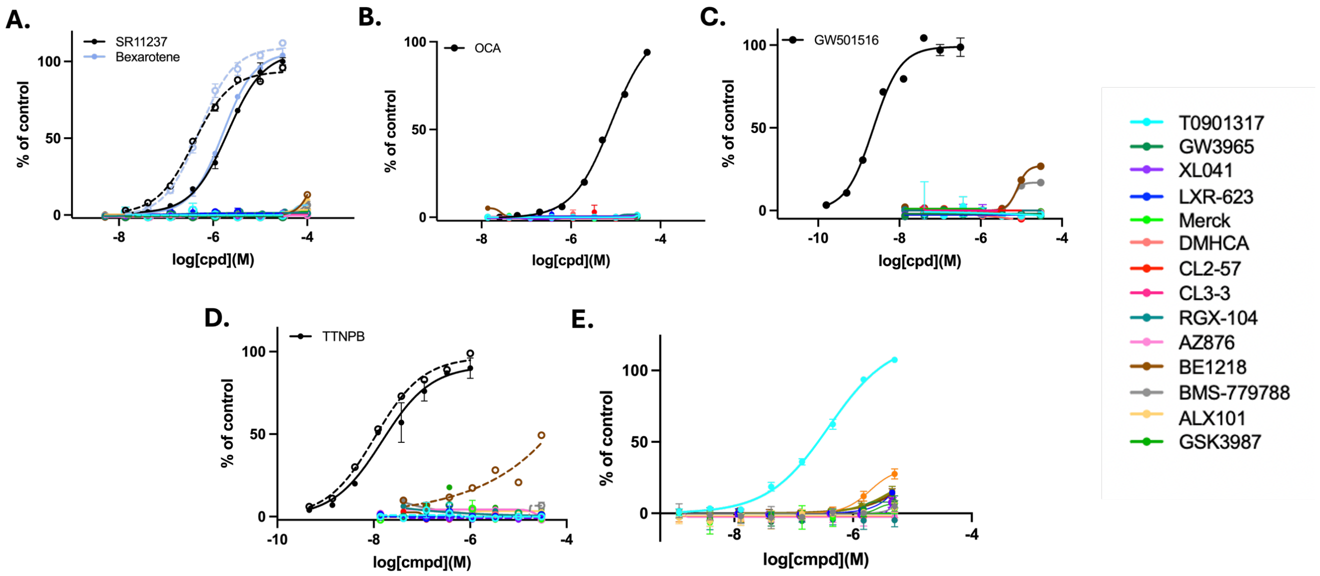

**Figure S1. NR Panel Results A.** Concentration-response for recruitment of coregulator D22 to RXRα (dotted lines) and RXRβ (solid line), determined by CRT assay, showing mean and SD, normalized to SR11237 as a full agonist. **B.** Concentration-response for recruitment of coregulator SRC2 to FXR determined by CRT assay, showing mean and SD, normalized to Obeticholic acid (OCA) as a full agonist. **C.** Concentration-response for recruitment of coregulator C33 to PPARδ determined by CRT assay, showing mean and SD, normalized to GW501516 as a full agonist. **D.** Concentration-response for recruitment of coregulator D22 to RARα (dotted lines) and recruitment of coregulator SRC2-2 RARβ (solid line), determined by CRT assay, showing mean and SD, normalized to TTNPB as a full agonist. **E.** Concentration-response for recruitment of coregulator SRC1-4 to PXR determined by CRT assay, showing mean and SD, normalized to T0901317(T0) as a full agonist.

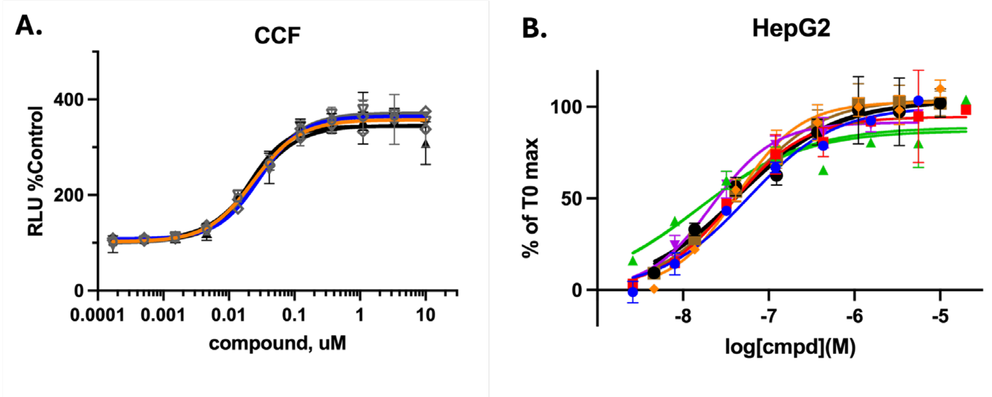

**Figure S2.** Concentration-response curves of the control compound, **T0**, in CCF ABCA1-luc reporter assay and HepG2 SRE-luc reporter assay (**A**, **B**) show high reproducibility across independent experiments.

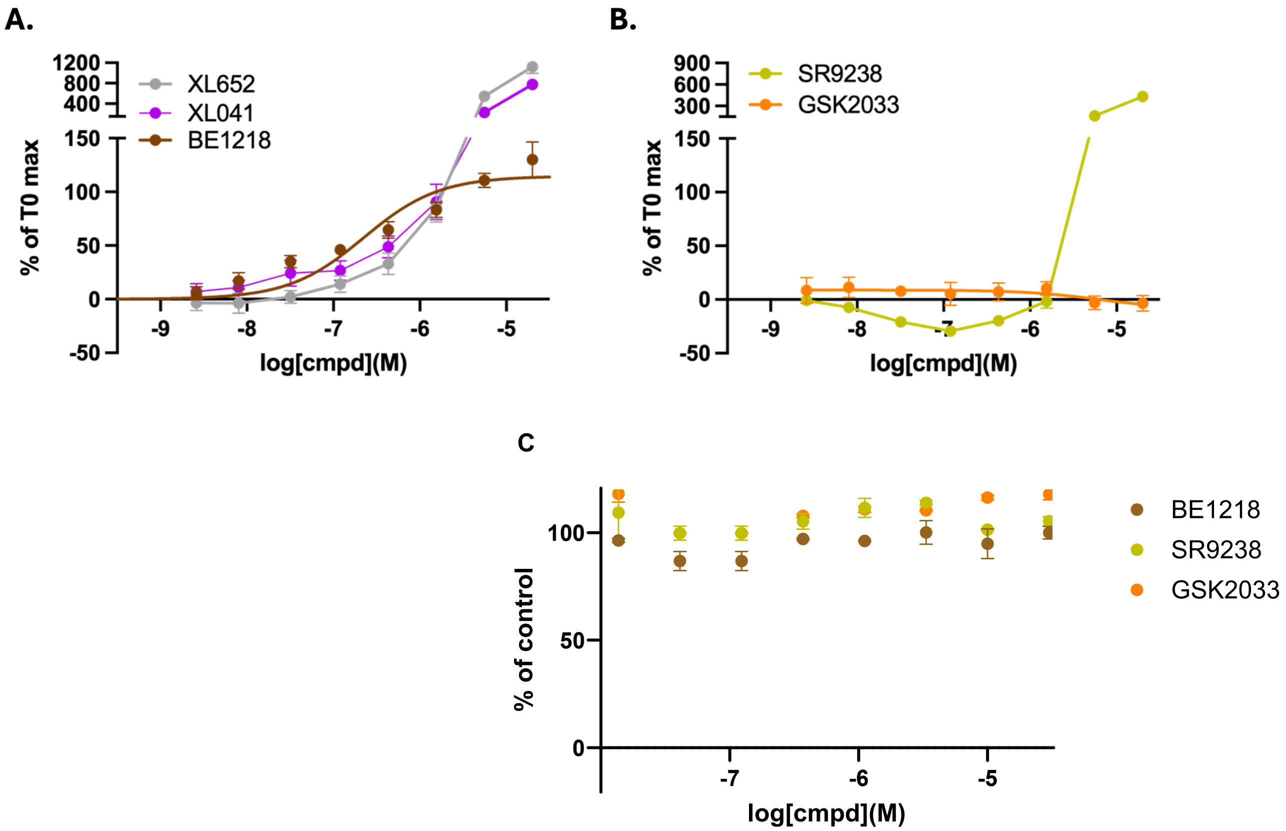

**Figure S3. A-B)** Lipogenic response of HepG2 cells to treatment with LXR ligands showing full concentration-response curves for biphenyl compounds shown in **Fig. 5E-F. C)** Cell viability in HepG2 cells under identical conditions to those used to obtain SRE-luc data.

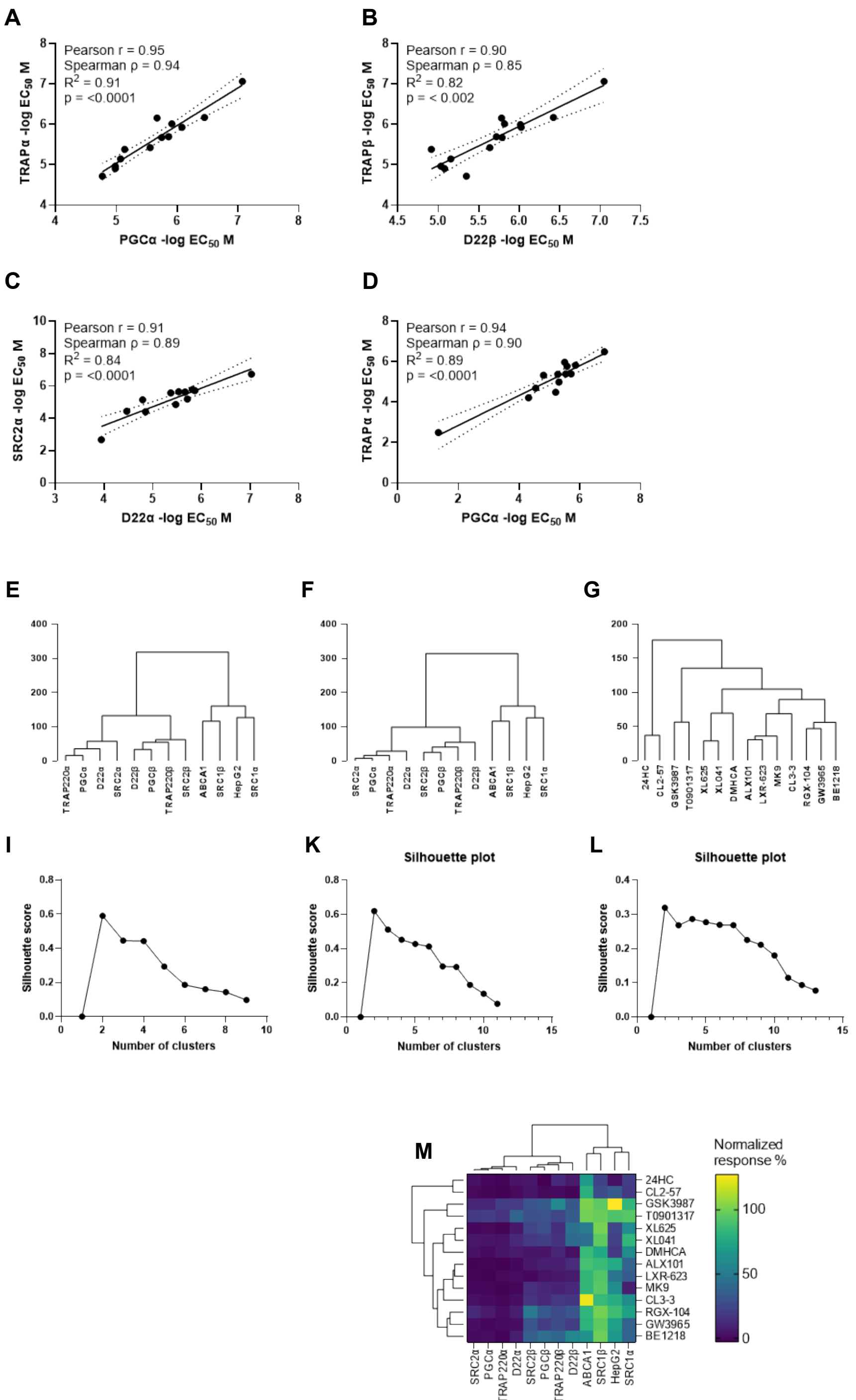

**Fig. S4. Correlating potency in cell-free and cell-based assays. A-D)** Correlation of potency for ligand-dependent recruitment of SRC2-2, D22, PGC1α, and TRAP220 to LXR α and β showing correlation statistics, best-fit line and 95% confidence intervals. Hierarchical clustering analysis of cell-free and cell-based relative response at 1 µM ligand (**E**) including all agonist ligands or (**F,G**) excluding AZ876. **I-L)** Silhouette plots for dendrograms **E, F, G,** respectively. **M)** Heatmap of hierarchical clustering for all ligands excluding AZ876. Raw data was analyzed in Prism 10.

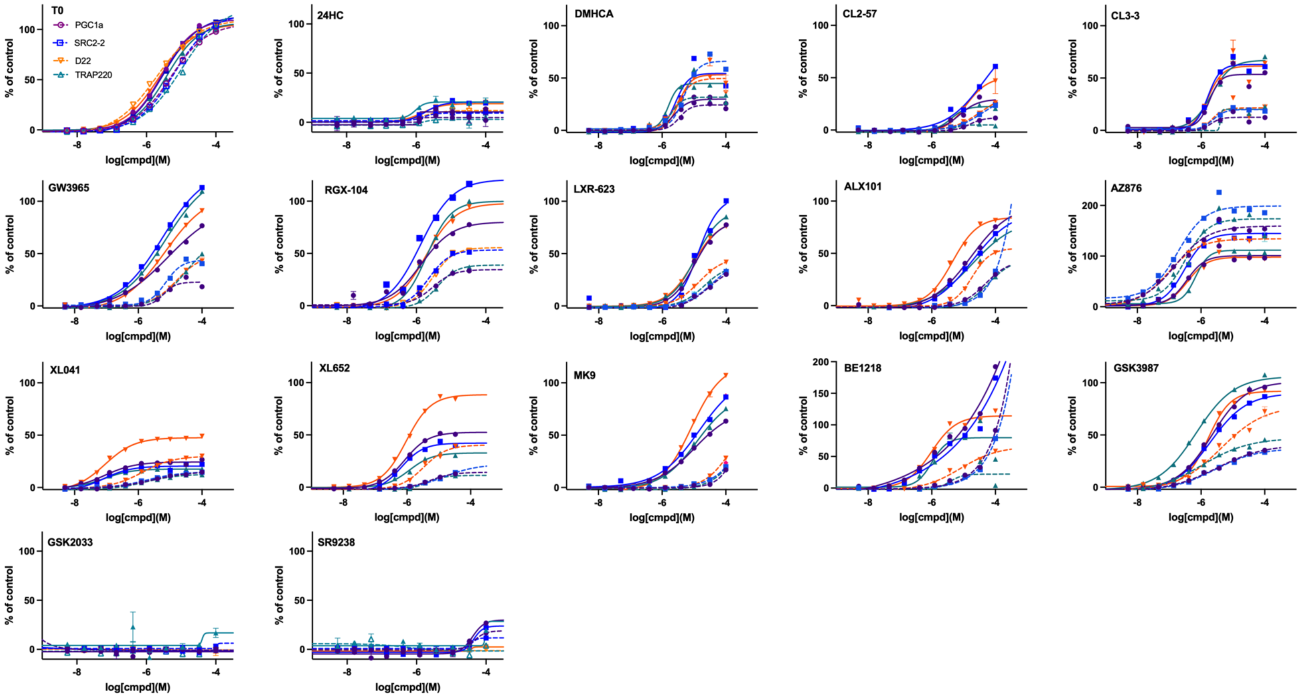

**Figure S5.** Full concentration-response profiles of all tested compounds across coactivators. Shown here for completeness; LXR$\alpha$ (dashed lines) and LXR$\beta$ (solid lines).

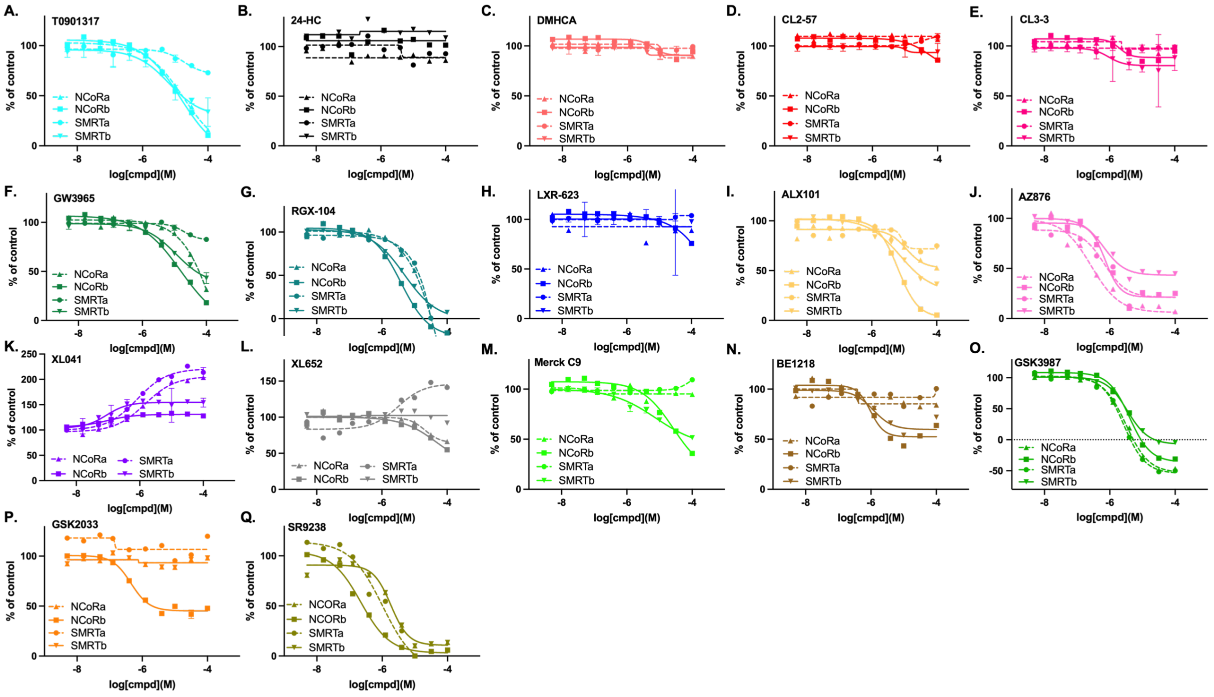

**Figure S6.** Full concentration-response profiles of all tested compounds across corepressors NCor2 and SMRT for both LXR$\alpha/\beta$ isoforms.

### Materials and Methods:

#### CRT Assays

**General statement about normalization:**

All controls were run on the same plate and on the same day, in parallel with the compound samples. Negative controls, DMSO, were used to define the 0% response, while 100% activity was defined by the maximal signal produced by the highest concentration of the appropriate positive control—T0901317 for LXR, and the corresponding full agonists for the other receptors.

**Coactivator recruitment LXRα/β HTRF**

To determine compound EC50 values, 10mM compound stocks were prepared in DMSO. An initial 8-concentration, 3-fold dilution series in DMSO was performed in 96-well v-bottom, polypropylene plates at 100x the final desired concentration. Intermediate dilution plates were prepared by diluting the 100x DMSO stocks 25-fold in assay buffer (50mM Tris, pH 7.4, 50mM KCl, 0.05% Triton-X 100, 10% glycerol, 0.1% BSA- (added fresh), 5mM DTT (added fresh)), creating a 4x stock. The following assay components were prepared individually at a 4x final concentration in assay buffer: (1) LXR$\alpha/\beta$ -GST tag (Thermo Fisher Scientific PV4657/PV PV4660); (2) a mixture of anti-GST-Tb (CisBio/Perkin Elmer 61GSTTLA, lot 14A) and SA-XL665 (CisBio/Perkin Elmer 610SAXLB); (3) SRC1-btn (AnaSpec AS62152/Fisher Scientific NC1565694). A 3uL aliquot of the 4x ligand was then added to each well of the 384-well small volume non-binding black assay plate (Greiner 784900) followed by a 3uL aliquot of each reaction component for a total volume of 12uL. Control wells used contained all components minus ligands, replaced by positive control T0901317 (Enzo 270-309-M010/VWR 89147-616) or DMSO. Incubated at room temperature for 1hr before measurement of TR-FRET. Final concentrations of assay components LXR$\alpha/\beta$ (4nM), anti-GST-Tb (0.4nM), SA-XL665(9.44nM), and SRC1-btn (100nM). TR-FRET measurements were performed on a CLARIOstar® Plus multi-mode plate reader (BMG Labtech, Ortenberg, Germany), excitation 337nm (laser), emission1 615nm and emission2 665nm. The signal is expressed as a ratio of TR-FRET measurements at 665 nm/615nm × 1000. Experimental data were normalized to controls and fit with a four parameter Hill equation using Graphpad Prism software (Boston, MA) to determine EC50s.

**Coactivator recruitment Lanthascreen RXRα/β**

To determine compound EC50 values, 10mM compound stocks were prepared in DMSO. An initial 8-concentration, 3-fold dilution series in DMSO was performed in 96-well v-bottom, polypropylene plates at 100x the final desired concentration. Intermediate dilution plates were prepared by diluting the 100x DMSO stocks 25-fold in assay buffer (50mM Tris, pH 7.4, 50mM KCl, 0.05% Triton-X 100, 10% glycerol, 0.1% BSA- (added fresh), 5mM DTT (added fresh)), creating a 4x stock. The following assay components were prepared individually at a 4x final concentration in assay buffer: (1) RXR$\alpha/\beta$ (Thermo Fisher Scientific PV4799/ PV4390); (2) anti-GST-Tb (CisBio/Perkin Elmer 61GSTTLA, lot 14A); (3) D22-FL (Thermo Fisher Scientific PV4386). A 3uL aliquot of the 4x ligand was then added to each well of the 384-well small volume non-binding black assay plate (Greiner 784900) followed by a 3uL aliquot of each reaction component for a total volume of 12uL. Control wells used contained all components minus ligands, replaced by positive control SR11273 (Sigma-Aldrich S8951) or DMSO. Incubated at room temperature for 1hr before measurement of TR-FRET. Final concentrations of assay components RXR$\alpha/\beta$ (1nM), anti-GST-Tb (1nM), and D22-FL (200nM). TR-FRET measurements were performed on a CLARIOstar® Plus multi-mode plate reader (BMG Labtech, Ortenberg, Germany), excitation 337nm (laser), emission1 520nm and emission2 490nm. The signal is expressed as a ratio of TR-FRET measurements at 520 nm/490nm × 1000. Experimental data were normalized to controls and fit with a four parameter Hill equation using Graphpad Prism software (Boston, MA) to determine EC50s.

**Coactivator recruitment Lanthascreen FXR**

To determine compound EC50 values, 10mM compound stocks were prepared in DMSO. An initial 8-concentration, 3-fold dilution series in DMSO was performed in 96-well v-bottom, polypropylene plates at 100x the final desired concentration. Intermediate dilution plates were prepared by diluting the 100x DMSO stocks 25-fold in assay buffer (50mM Tris, pH 7.4, 50mM KCl, 0.05% Triton-X 100, 10% glycerol, 0.1% BSA- (added fresh), 5mM DTT (added fresh)), creating a 4x stock. The following assay components were prepared individually at a 4x final concentration in assay buffer: (1) FXR (Thermo Fisher Scientific **PV4835)**; (2) anti-GST-Tb (CisBio/Perkin Elmer 61GSTTLA, lot 14A); (3) SRC2-2-FL (Thermo Fisher Scientific PV4586). A 3uL aliquot of the 4x ligand was then added to each well of the 384-well small volume non-binding black assay plate (Greiner 784900) followed by a 3uL aliquot of each reaction component for a total volume of 12uL. Control wells used contained all components minus ligands, replaced by positive control Obeticholic acid (Sigma-Aldrich SML3096) or DMSO. Incubated at room temperature for 1hr before measurement of TR-FRET. Final concentrations of assay components FXR (2.5nM), anti-GST-Tb (1nM), and SRC2-2-FL (250nM). TR-FRET measurements were performed on a CLARIOstar® Plus multi-mode plate reader (BMG Labtech, Ortenberg, Germany), excitation 337nm (laser), emission1 520nm and emission2 490nm. The signal is expressed as a ratio of TR-FRET measurements at 520 nm/490nm × 1000. Experimental data were normalized to controls and fit with a four parameter Hill equation using Graphpad Prism software (Boston, MA) to determine EC50s.

**Coactivator recruitment Lanthascreen PPARδ**

To determine compound EC50 values, 10mM compound stocks were prepared in DMSO. An initial 8-concentration, 3-fold dilution series in DMSO was performed in 96-well v-bottom, polypropylene plates at 100x the final desired concentration. Intermediate dilution plates were prepared by diluting the 100x DMSO stocks 25-fold in assay buffer ((20mM Tris, pH 7.9, 100mM KCl, 0.01% Triton-X 100, 0.1% BSA (added fresh)), creating a 4x stock. The following assay components were prepared individually at a 4x final concentration in assay buffer: (1)PPAR$\delta$ (Thermo Fisher Scientific PV4694**)**; (2) anti-GST-Tb (CisBio/Perkin Elmer 61GSTTLA, lot 14A); (3) C33-FL (Thermo Fisher Scientific PV4606). A 3uL aliquot of the 4x ligand was then added to each well of the 384-well small volume non-binding black assay plate (Greiner 784900) followed by a 3uL aliquot of each reaction component for a total volume of 12uL. Control wells used contained all components minus ligands, replaced by positive control GW501516 (MedChemExpress, HY-10838/CS-0438) or DMSO. Incubated at room temperature for 1hr before measurement of TR-FRET. Final concentrations of assay components PPAR$\delta$ (5nM), anti-GST-Tb (1nM), and C33-FL (250nM). TR-FRET measurements were performed on a CLARIOstar® Plus multi-mode plate reader (BMG Labtech, Ortenberg, Germany), excitation 337nm (laser), emission1 520nm and emission2 490nm. The signal is expressed as a ratio of TR-FRET measurements at 520 nm/490nm × 1000. Experimental data were normalized to controls and fit with a four parameter Hill equation using Graphpad Prism software (Boston, MA) to determine EC50s.

**Coactivator recruitment Lanthascreen RARα.**

To determine compound $EC_{50}$ values, 10mM compound stocks were prepared in DMSO. An initial 8-concentration, 3-fold dilution series in DMSO was performed in 96-well v-bottom, polypropylene plates at 100x the final desired concentration. Intermediate dilution plates were prepared by diluting the 100x DMSO stocks 25-fold in assay buffer (50 mM Tris, pH 7.4, 50 mM KCl, 0.05% Triton-X 100, 10% glycerol, 0.1% BSA- (added fresh), 5 mM DTT (added fresh)), creating a 4x stock. The following assay components were prepared individually at a 4x final concentration in assay buffer: (1) RAR$\alpha$ (Thermo Fisher Scientific PV4799/ PV4390); (2) anti-GST-Tb (CisBio/Perkin Elmer 61GSTTLA, lot 14A); (3) D22-FL (Thermo Fisher Scientific PV4386). A 3 µL aliquot of the 4x ligand was then added to each well of the 384-well small volume non-binding black assay plate (Greiner 784900) followed by a 3 µL aliquot of each reaction component for a total volume of 12 µL. Control wells used contained all components minus ligands, replaced by positive control TTNPB (MedChemExpress HY-15682), or DMSO. Incubated at room temperature for 1hr before measurement of TR-FRET. Final concentrations of assay components RAR$\alpha$ (3 nM), anti-GST-Tb (2 nM), and D22-FL (250 nM). TR-FRET measurements were performed on a CLARIOstar® Plus multi-mode plate reader (BMG Labtech, Ortenberg, Germany), excitation 337nm (laser), emission1 520 nm and emission2 490 nm. The signal is expressed as a ratio of TR-FRET measurements at 520 nm/490 nm × 1000. Experimental data were normalized to controls and fit with a four parameter Hill equation using Graphpad Prism software (Boston, MA) to determine $EC_{50}$ values.

**Coactivator recruitment Lanthascreen RARβ.**

To determine compound $EC_{50}$ values, 10mM compound stocks were prepared in DMSO. An initial 8-concentration, 3-fold dilution series in DMSO was performed in 96-well v-bottom, polypropylene plates at 100x the final desired concentration. Intermediate dilution plates were prepared by diluting the 100x DMSO stocks 25-fold in assay buffer (50 mM Tris, pH 7.4, 50 mM KCl, 0.05% Triton-X 100, 10% glycerol, 0.1% BSA- (added fresh), 5 mM DTT (added fresh)), creating a 4x stock. The following assay components were prepared individually at a 4x final concentration in assay buffer: (1) RAR$\beta$ (Thermo Fisher Scientific PV4799/ PV4390); (2) anti-GST-Tb (CisBio/Perkin Elmer 61GSTTLA, lot 14A); (3) SRC2-2-FL (Thermo Fisher Scientific PV4586). A 3 µL aliquot of the 4x ligand was then added to each well of the 384-well small volume non-binding black assay plate (Greiner 784900) followed by a 3 µL aliquot of each reaction component for a total volume of 12 µL. Control wells used contained all components minus ligands, replaced by positive control TTNPB (MedChemExpress HY-15682) or DMSO. Incubated at room temperature for 1hr before measurement of TR-FRET. Final concentrations of assay components RAR$\beta$ (2.5 nM), anti-GST-Tb (2 nM), and SRC2-2-FL (300 nM). TR-FRET measurements were performed on a CLARIOstar® Plus multi-mode plate reader (BMG Labtech, Ortenberg, Germany), excitation 337 nm (laser), emission1 520 nm and emission2 490 nm. The signal is expressed as a ratio of TR-FRET measurements at 520 nm/490 nm × 1000. Experimental data were normalized to controls and fit with a four parameter Hill equation using Graphpad Prism software (Boston, MA) to determine $EC_{50}$ values.

**Coactivator recruitment Lanthascreen PXR.** To determine compound $EC_{50}$ values, 10 mM compound stocks were prepared in DMSO. An initial 8-concentration, 3-fold dilution series in DMSO was performed in 96-well v-bottom, polypropylene plates at 100x the final desired concentration. Intermediate dilution plates were prepared by diluting the 100x DMSO stocks 33.3-fold in assay buffer (50 mM Hepes, pH 8.0, 50 mM NaCl, 0.01% TWEEN® 20), creating a 3x stock. The following assay components were prepared individually at a 3x final concentration in assay buffer: (1) PXR (Thermo Fisher Scientific PV4841); (2) anti-GST-Tb (Revvity 61GSTTLA) combined with SRC1-4-FL (Thermo Fisher Scientific PV4582). A 4 µL aliquot of the 3x ligand was then added to each well of the 384-well small volume non-binding black assay plate (Greiner 784900) followed by a 4 µL aliquot of each reaction component for a total volume of 12 µL. Control wells used contained all components minus ligands, replaced by positive control T0901317 (Synthesized by Ganga Reddy Velma, Thatcher Lab, University of Arizona; or Enzo 270-309-M010/VWR 89147-616) or DMSO. Final concentrations of assay components were: PXR (10 nM), anti-GST-Tb (1 nM), and SRC1-4-FL (300 nM). Plates were incubated for 20' at room temperature before TR-FRET measurements were performed on a CLARIOstar® Plus multi-mode plate reader (BMG Labtech, Ortenberg, Germany), excitation 337 nm (laser), emission1 520 nm and emission2 490 nm. The signal is expressed as a ratio of TR-FRET measurements at 520 nm/490 nm × 1000. Experimental data were normalized to controls and fit with a four parameter Hill equation using Graphpad Prism software (Boston, MA) to determine $EC_{50}$ values.

**Chemistry**

(*R*)-2-chloro-4-(1'-(2-hydroxy-3-methyl-2-(trifluoromethyl)butanoyl)-[4,4'-bipiperidin]-1-yl)-*N*,*N*-dimethylbenzamide (**MK9**)

This compound was obtained using a procedure similar to the reported literature method.[1]

$^{1}$H NMR (400 MHz, $CDCl_3$) $\delta$ 7.14 (d, $J$ = 8.5 Hz, 1H), 6.85 (d, $J$ = 2.4 Hz, 1H), 6.80 (dd, $J$ = 8.6, 2.4 Hz, 1H), 5.69 (s, 1H), 4.88 – 4.14 (m, 2H), 3.74 (dd, $J$ = 12.7, 4.0 Hz, 2H), 3.11 (s, 3H), 2.88 (s, 3H), 2.71 (td, $J$ = 12.1, 2.3 Hz, 2H), 2.44 (s, 1H), 1.90 – 1.76 (m, 4H), 1.45 – 1.16 (m, 6H), 1.11 (d, $J$ = 6.4 Hz, 3H), 0.84 (s, 3H). $^{13}$C NMR (101 MHz, $CDCl_3$) $\delta$ 169.14, 166.93, 152.49, 131.42, 128.74, 126.13, 123.17, 115.98, 114.44, 49.23, 46.49, 40.96, 40.58, 38.43, 34.89, 31.07, 29.48, 28.92, 28.87, 16.78.

*Ethyl3-(5-chloro-3-(N-(3,4-diethoxyphenyl)-N-methylsulfamoyl)thiophene-2-carboxamido)benzoate* (**CL2-27)**

This compound was obtained using procedures detailed in our previous publication.[2]

$^{1}$H NMR (400 MHz, $CDCl_3$) δ 10.03 (s, 1H), 7.94 (t, $J$ = 1.9 Hz, 1H), 7.77 (dt, $J$ = 7.8, 1.3 Hz, 1H), 7.59 (ddd, $J$ = 8.2, 2.3, 1.1 Hz, 1H), 7.31 (t, $J$ = 7.9 Hz, 1H), 6.63 – 6.57 (m, 2H), 6.53 (d, $J$ = 8.5 Hz, 1H), 4.39 (q, $J$ = 7.1 Hz, 2H), 3.87 (q, $J$ = 6.9 Hz, 2H), 3.70 (q, $J$ = 6.9 Hz, 2H), 3.20 (s, 3H), 1.41 (t, $J$ = 7.1 Hz, 3H), 1.34 (t, $J$ = 7.0 Hz, 3H), 1.26 (t, $J$ = 7.0 Hz, 3H). $^{13}$C NMR (101 MHz, $CDCl_3$) δ 166.00, 156.54, 149.21, 149.10, 142.56, 137.44, 135.28, 132.04, 131.83, 131.13, 130.13, 128.71, 125.64, 123.72, 120.37, 119.12, 112.73, 112.54, 77.34, 77.02, 76.71, 64.65, 64.37, 61.12, 38.89, 14.70, 14.55, 14.37.

*Ethyl3-chloro-5-(5-chloro-3-(N-(4-ethoxy-3-methoxyphenyl)-Nmethylsulfamoyl) thiophene-2-carboxamido)benzoate* (**CL3-3**)

This compound was obtained using procedures detailed in our previous publication.[2]

$^{1}$H NMR (400 MHz, $CDCl_3$) δ 10.00 (s, 1H), 7.72 (p, $J$ = 2.0 Hz, 2H), 7.68 (t, $J$ = 1.8 Hz, 1H), 7.34 (s, 1H), 6.66 (d, $J$ = 2.1 Hz, 1H), 6.56 – 6.53 (m, 2H), 4.38 (q, $J$ = 7.1 Hz, 2H), 3.75 (q, $J$ = 7.0 Hz, 2H), 3.71 (s, 3H), 1.41 (t, $J$ = 7.1 Hz, 3H), 1.31 (t, $J$ = 7.0 Hz, 3H). $^{13}$C

NMR (101 MHz, $CDCl_3$) δ 164.86, 156.70, 149.58, 148.95, 141.95, 138.42, 135.73, 134.70, 132.33, 132.00, 131.94, 130.24, 125.37, 123.18, 118.67, 118.29, 111.80, 111.14, 77.34, 77.02, 76.70, 64.24, 61.54, 55.91, 38.87, 14.51, 14.31.

### Spectra

#### $^{1}$H NMR of **MK9**

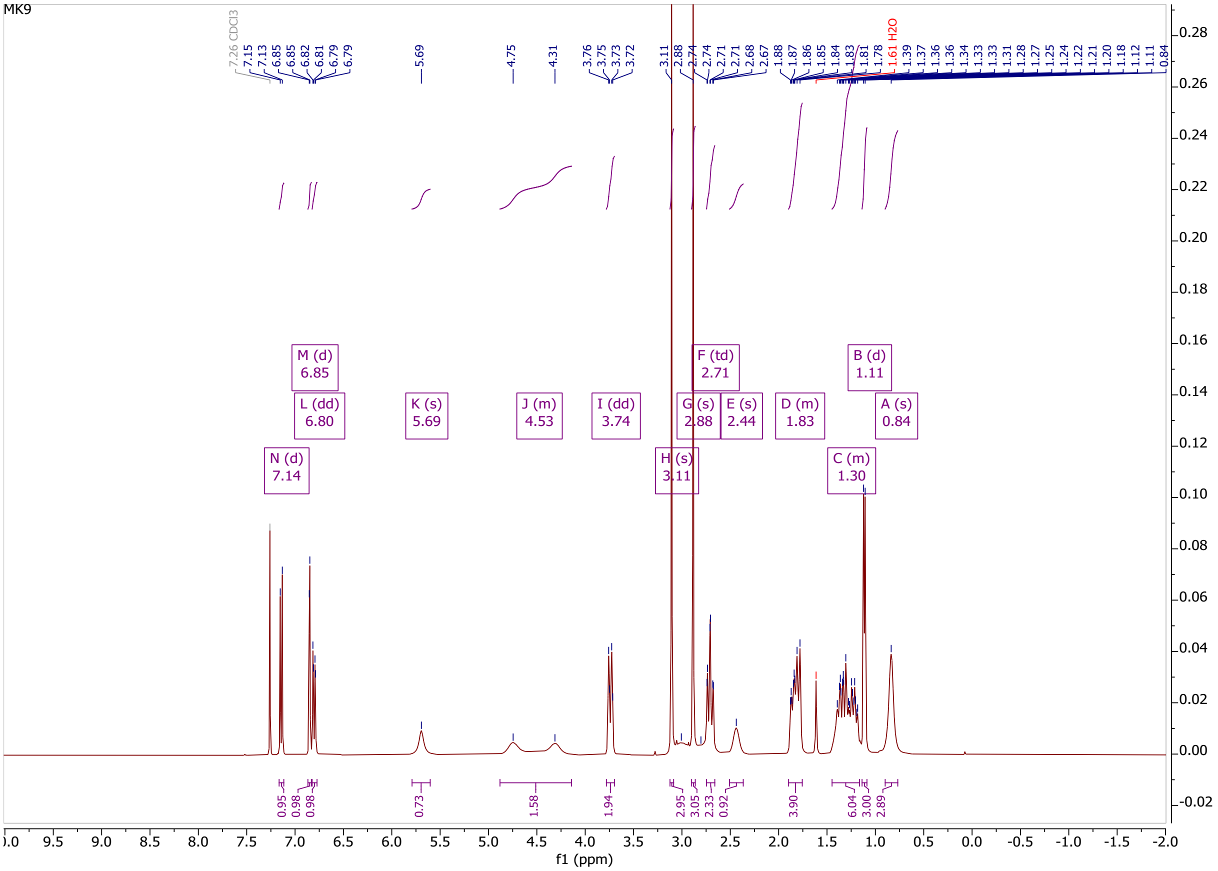

$^{13}C$ NMR of **MK9**

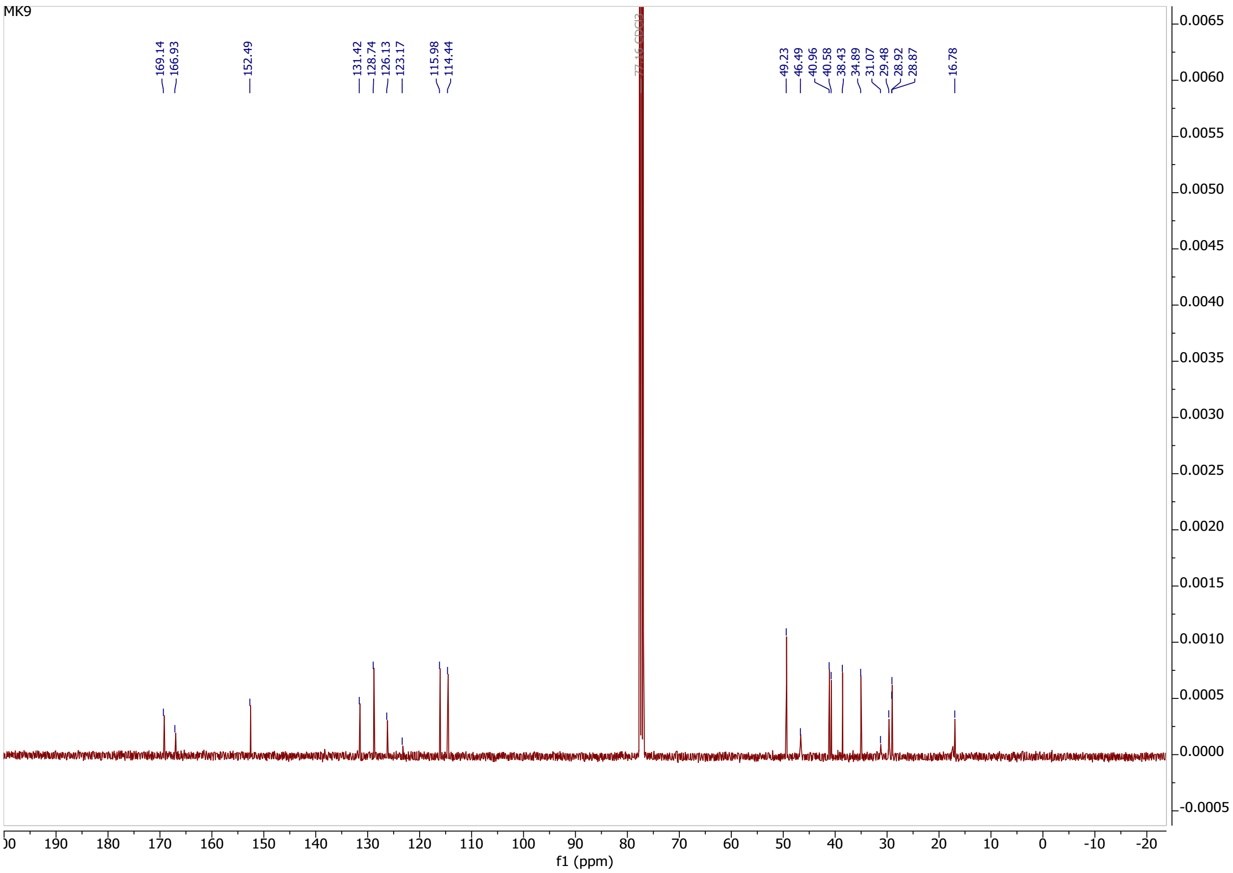

[1]H NMR of **CL2-57**

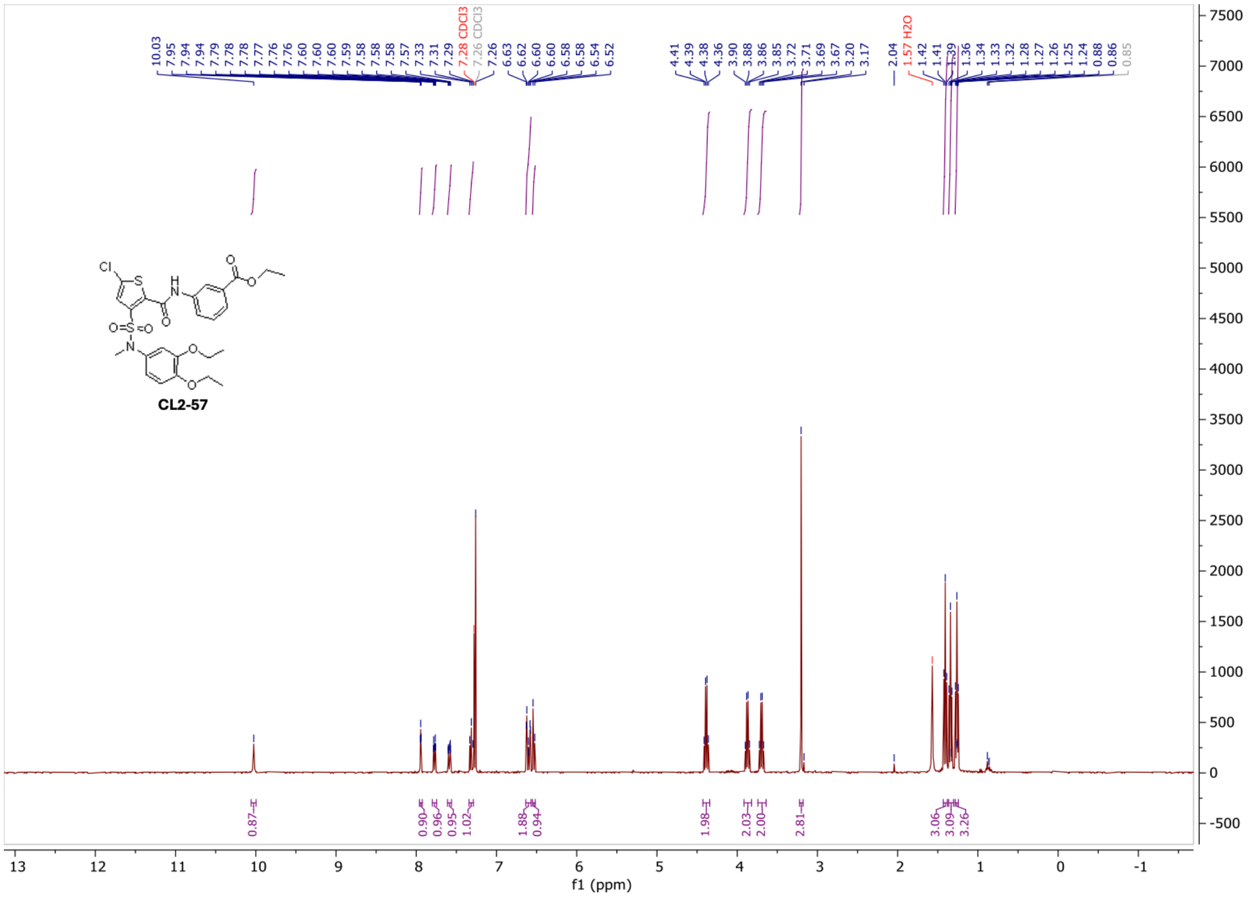

$^{13}$C NMR of **CL2-57**

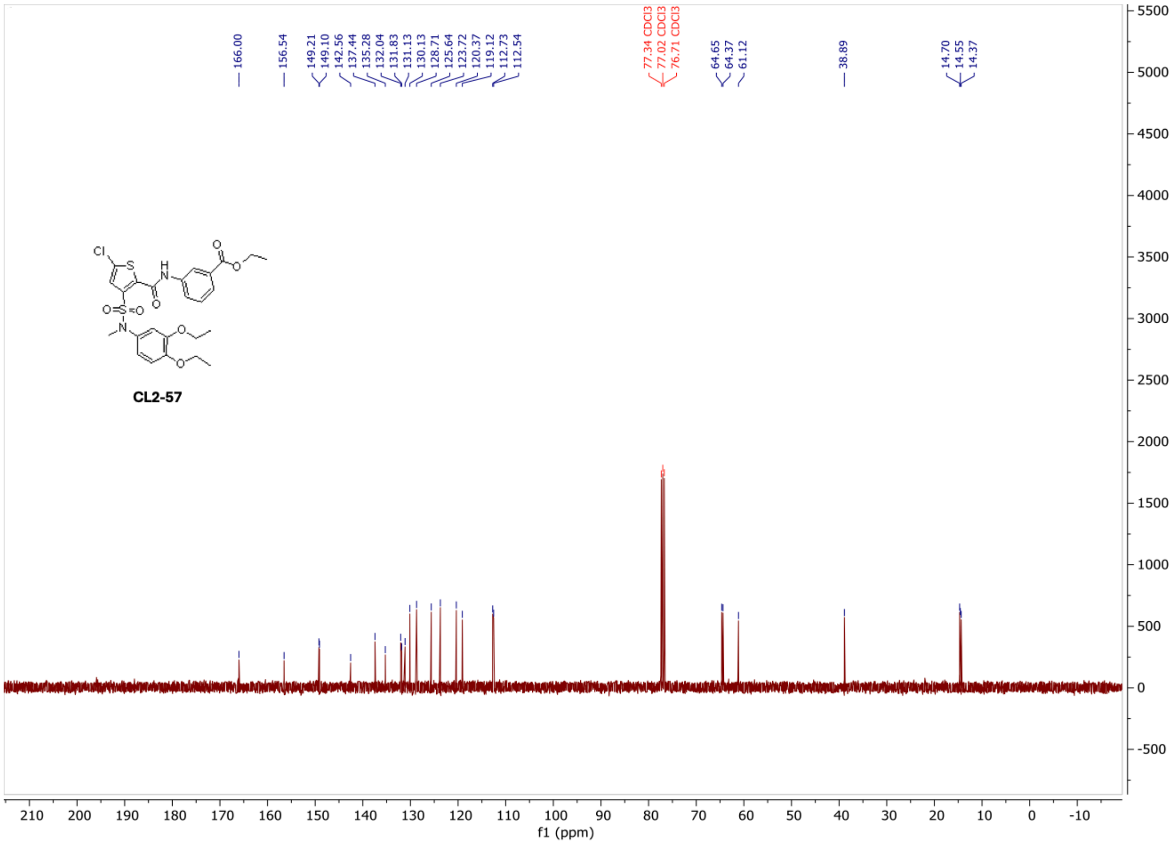

[1]H NMR of **CL3-3**

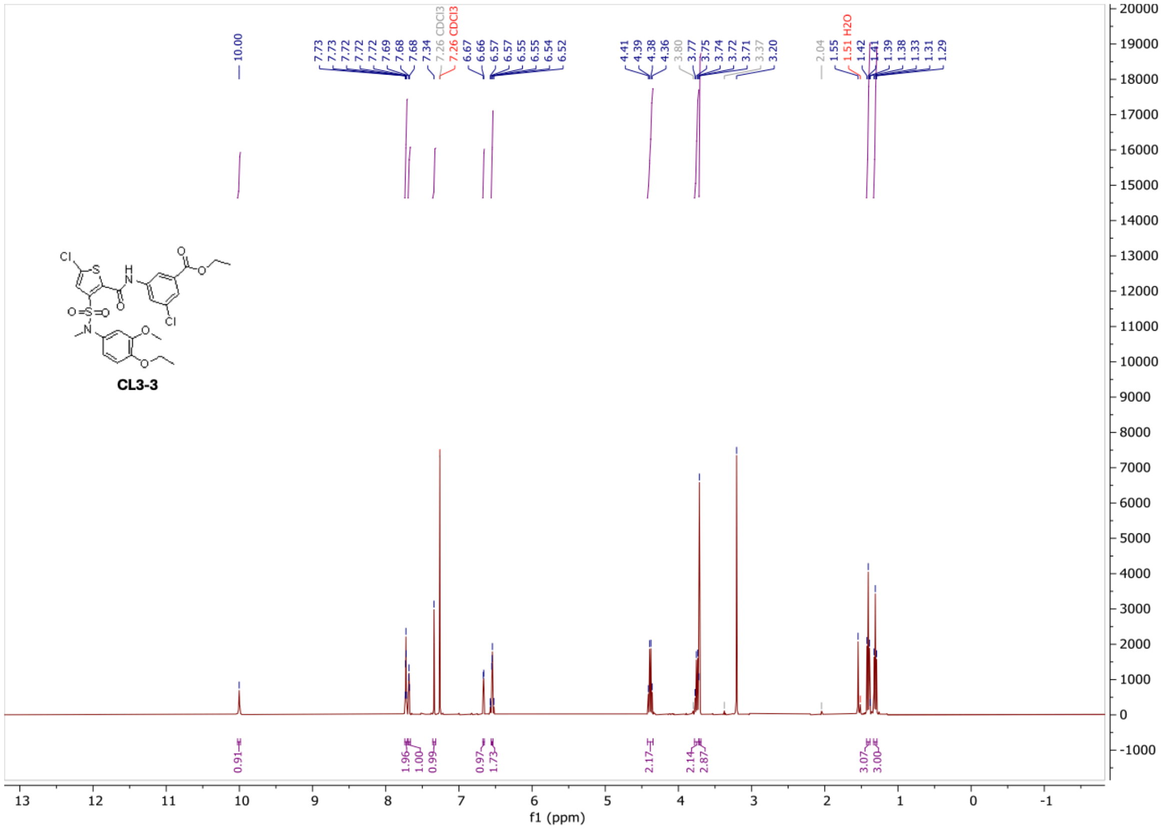

$^{13}$C NMR of **CL3-3**

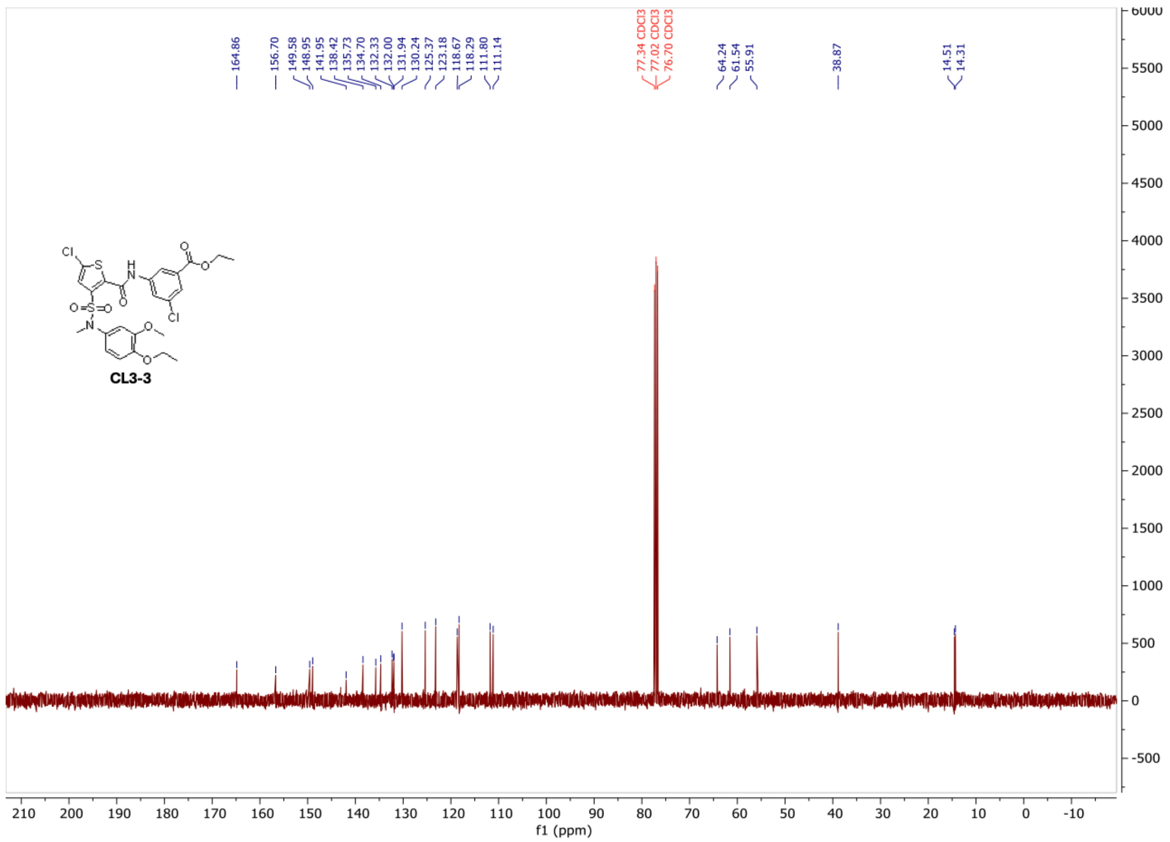

(1) Stachel, S. J.; Zerbinatti, C.; Rudd, M. T.; Cosden, M.; Suon, S.; Nanda, K. K.; Wessner, K.; DiMuzio, J.; Maxwell, J.; Wu, Z.; et al. Identification and in Vivo Evaluation of Liver X Receptor β-Selective Agonists for the Potential Treatment of Alzheimer's Disease. *Journal of Medicinal Chemistry* **2016**, *59* (7), 3489-3498. DOI: 10.1021/acs.jmedchem.6b00176.
(2) Velma, G. R.; Laham, M. S.; Lewandowski, C.; Valencia-Olvera, A. C.; Balu, D.; Moore, A.; Ackerman-Berrier, M.; Rychetsky, P.; Penton, C.; Musku, S. R.; et al. Nonlipogenic ABCA1 Inducers (NLAI) for Alzheimer's Disease Validated in a Mouse Model Expressing Human APOE3/APOE4. *Journal of Medicinal Chemistry* **2024**, *67* (17), 15061-15079. DOI: 10.1021/acs.jmedchem.4c00733.
